# A Robust Computational Framework for the Optimization of CDK7 Inhibitors as Promising Cancer Therapy

**DOI:** 10.1101/2024.11.13.623385

**Authors:** Zhaoqi Shi, Xufan Gao, Damiano Buratto, Ruhong Zhou

## Abstract

Cyclin-dependent kinase 7 (CDK7) plays a crucial role in cell cycle regulation and transcription, establishing it as a promising target for cancer therapy. Although the covalent inhibitor THZ1 effectively targets CDK7, it presents risks such as short half-life and potential off-target side effects. To address these limitations, we employed a computational workflow integrating virtual screening, molecular dynamics (MD) simulations, and free energy perturbation (FEP) method to design non-covalent CDK7 inhibitors with enhanced selectivity and safety profiles. MD simulations elucidated THZ1’s inhibitory mechanism and identified key molecular fragments within its structure. By incorporating fragments from known inhibitors, we introduced extensive non-covalent interactions within the binding pocket, leading to the identification of three novel non-covalent inhibitors with binding affinities comparable to or higher than that of THZ1. Our findings not only introduce promising CDK7 inhibitors but also present a robust computational framework that could accelerate the discovery of kinase-targeted therapeutics.

## 1 Introduction

Cycle-dependent kinase 7 (CDK7) forms a complex with Cyclin H and MAT1 to create the CDK activating kinase (CAK) complex^1^. The CAK complex performs various biological functions, primarily regulating gene transcription and driving the normal cell cycle progression. As part of transcription factor IIH (TFIIH), CDK7 phosphorylates the Ser5 in the C-terminal domain (CTD) of RNA polymerase II (RNA Pol II) (Figure 1A)^2^. Additionally, CDK7 phosphorylates CDK9, which further phosphorylates the Ser2 of RNA-Pol II CTD^3^. These phosphorylation events are essential to facilitate transcription elongation. CDK7 activity is negatively regulated by the XPD subunit in TFIIH, which impedes transcription and the mitotic cell cycle^4^. Within the cell cycle, CDK7 activates various other CDKs at different stages (Figure 1B)^5^. Inhibiting CDK7 during the G1 phase blocks CDK2 activation and delays the entry into the S phase. CDK7 also phosphorylates the T-loop of CDK4/6^6^, thus affecting the G1-to-S phase transition. Meanwhile, inhibiting CDK7 in the G2 phase disrupts the assembly of the CDK1/Cyclin B complex, preventing cells from entering mitosis^7^. Given its critical roles in transcriptional regulation and cell cycle progression, CDK7 has emerged as a promising therapeutic target in cancers^1,8^. A substantial body of evidence indicates that CDK7 is crucial in maintaining the malignant phenotype of various cancer cells and its overexpression has been observed in various tumors, making it a potential prognostic marker^9^. For instance, CDK7 overexpression in triple-negative breast cancer (TNBC) correlates with increased cell proliferation and survival, while CDK7 inhibition leads to the apoptosis in TNBC cells^10^. CDK7 inhibitors can also effectively control the proliferation and survival of lung squamous cell carcinoma (LSCC) cells by reducing amplified SOX2 expression^11^. Moreover, CDK7 is found to be critically involved in the transcriptional regulation of undifferentiated thyroid carcinoma (ATC)^12^. These findings underscore the potential of CDK7 inhibitors in cancer treatment.

**Figure 1.**
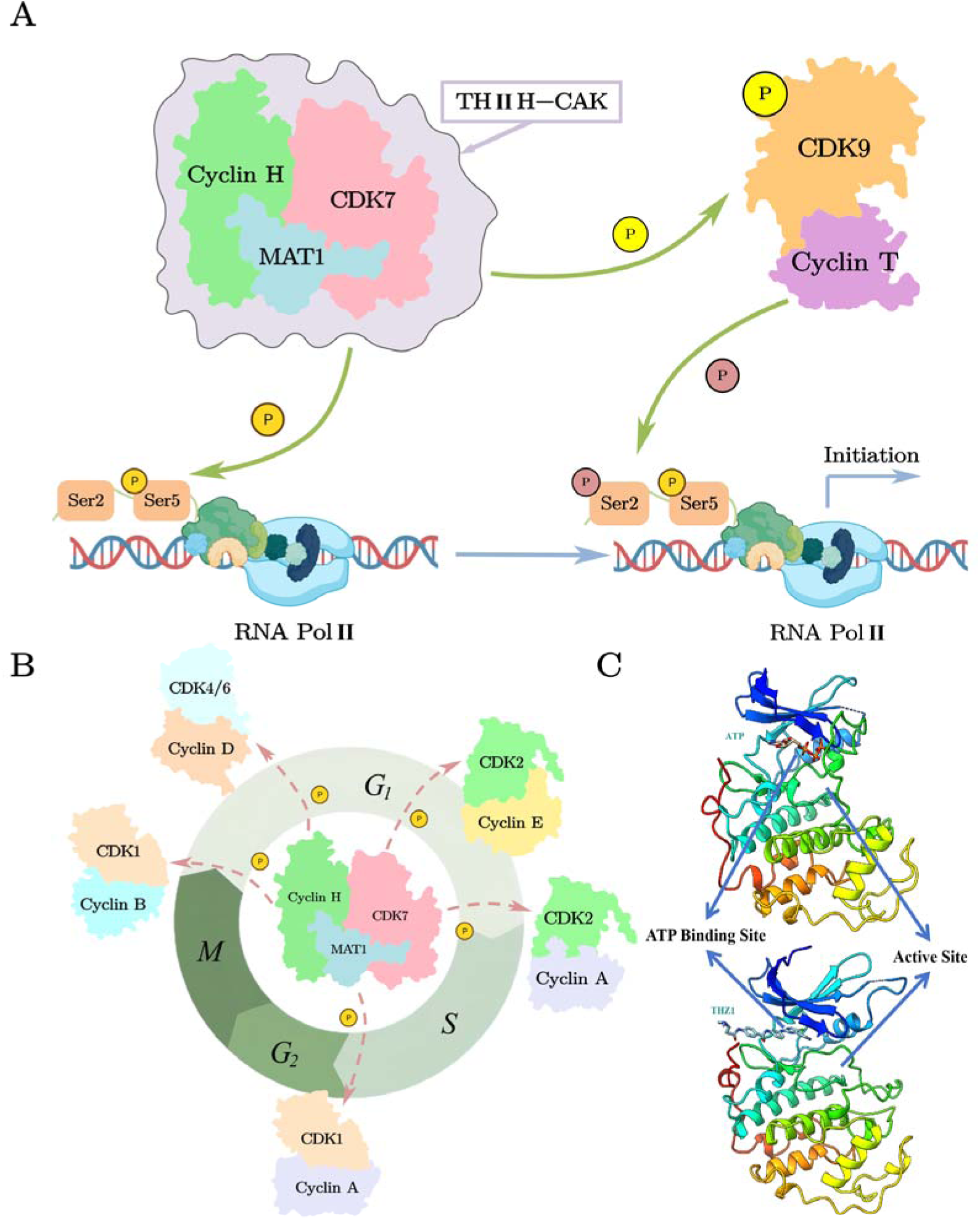
Biological functions and structural features of CDK7. Circles labeled with “P” denote phosphate groups. (A) Role in transcription: As part of the TFIIH complex, CDK7 promotes transcription by specifically phosphorylating RNA polymerase II at Ser5. It also facilitates transcription elongation through CDK9 activation. (B) Role in the cell cycle: CDK7 regulates the cell cycle by activating CDK1 and CDK2 at different phases and phosphorylating the T-loop of CDK4/6, thereby driving cell cycle progression. (C) Structural features: The ATP-binding site and the active sites of CDK7 are shown, highlighting critical regions targeted by inhibitors for enhanced specificity and effectiveness.

Approaches for developing CDK7 inhibitors include non-selective CDK inhibitors (pan CDK inhibitors) and selective CDK7 inhibitors, the latter of which are further divided into non-covalent inhibitors and irreversible covalent inhibitors^8,13,14^. Early non-selective pan CDK inhibitors like Flavopiridol^15^, SYS-032^16^, and Roscovitine^17^ faced challenges in clinical settings due to the structural similarity of CDK kinase domains^14^. BS-181, the first selective non-covalent CDK7 inhibitor^18^, led to further developments, including LDC4297, which mediates the downregulation of p53 expression in TNBC cells^19^, YKL-5-124, which blocks the cell cycle and activates the immune responses^20^, and SY-5609, which is currently in phase I clinical trials^21^. On the other hand, covalent inhibitors can form bonds with the target proteins, potentially offering more durable and effective inhibition with improved selectivity^22^. Recently, more attention has been given to the covalent inhibitor THZ1 (Figure 1C)^23^, which binds covalently to Cys312 of CDK7 via a Michael addition reaction with its α,β-unsaturated ketone moiety, while partially blocking the ATP-binding pocket through stable non-covalent interactions. THZ1 showed strong inhibitory potency against CDK7 (with an IC50 of 3.2 nM^24^), and demonstrated antitumor efficacy across multiple cancer types such as TNBC, neuroblastoma and castration-resistant prostate cancer (CRPC)^25^, urothelial carcinoma^26^, colorectal cancer^27^, chordoma^28^ and non-small cell lung cancer (NSCLC)^29^. However, THZ1’s short plasma half-life (∼45 minutes) limits its therapeutic duration, prompting the development of derivatives such as THZ2, which extends the half-life by fivefold while maintaining the specificity^10^. Moreover, resistance to THZ1 can develop through upregulation of ABC transporters (e.g. ABCB1), reducing intracellular drug accumulation and, consequently, its efficacy in cancer cells^30^. Additionally, off-target effects on CDK12 and CDK13 have been observed. While these effects may strengthen the suppression of super-enhancer-associated genes, they raise concerns about unintended impacts on non-cancerous cells^31^. The high degree of homology within the CDK family^32^ presents further challenges, as inhibitors targeting only the ATP-binding site may cause off-target side effects^33^. Consequently, non-covalent CDK7 inhibitors offers a promising alternative for improved safety and reduced side effects.

To address current CDK7 inhibitors limitations, this study employs a series of computational techniques to design non-covalent CDK7 inhibitors with enhanced selectivity and minimized off-target effects. Given the shared core structure of THZ1 and THZ2^34^, which is essential for binding with CDK7, the interaction of this core segment with CDK7 has been examined in atomic detail, along with its role in stabilizing THZ1. In this study, we considered both the ATP-binding site and the active site of CDK7 in our virtual screening process (Figure 1C) to potentially enhance selectivity over other CDKs^35^. To further quantify the binding affinity of our designed inhibitors, we utilized free energy perturbation (FEP), which are known for their accuracy in calculating binding free energy differences between a ligand and its analogue^36^. Also, FEP methods have been demonstrated to be highly consistent with experimental results^37^ and have been successfully applied in drug discovery and vaccine design, despite their computational intensity^38^. With molecular dynamics simulations and FEP calculations, we identified three stable small-molecule inhibitors with high affinity for CDK7 without relying on covalent bonds. This work not only presents safer alternatives to covalent CDK7 inhibitors but also demonstrates a versatile computational framework for drug discovery, offering valuable insights for kinase-targeted therapies.

## 2 Materials and Methods

### 2.1 System Preparation

The structure of the human CAK (CDK7, Cyclin H and MAT1) in complex with covalent bounded THZ1 was obtained from a Cryo-EM structure (PDB ID: 6XD3)^39^. Phosphorylation of Thr170 and Ser164 is reported to enhance the kinase activity, strengthen Cyclin H binding, and influence CAK substrate specificity^1,40^. These phosphorylation sites were incorporated into our model and SWISS-MODEL^41^ was used to add the missing residues. All histidine residues were protonated only at the Nε atom. In the text, we refer to the structure of CDK7 in complex with covalent bound THZ1 as THZ1-Cov. Moreover, to explain the binding mechanism of THZ1 and identify its core segment, three more systems were built: THZ1-Noncov, THZ1-Segment1 and THZ1-Segment2 (Figure 3A). THZ1-Noncov was obtained from THZ1-Cov by removing the covalent bond, simulating the occasion of not undergoing Michael addition between the THZ1 and CDK7. THZ1-Segment1 and THZ1-Segment2 were both obtained from of THZ1-Noncov by cutting part of the molecule (Figure 3A).

### 2.2 Virtual Screening

To identify potential CDK7 inhibitors, multi-step virtual screening approach were applied, as outlined in Figure 2:

1. Similarity search. Eight known CDK7 small molecule inhibitors have been reported in the literature^8^ and 147 molecules were collected from PubChem database with a similarity to these known inhibitors above 97% ^42^.
2. First round of molecular docking. The 147 molecules were docked to the CDK7 binding pocket using smina^43,44^. The docking box was set to include pocket 1, pocket 2, and the core pocket (Figure 3G), where the core pocket is the main binding site for THZ1-Segment2. This step generated plausible conformations of the molecules in complex with the receptor.
3. Fragment extraction. We extracted molecule fragments bound to CDK7 in pocket 1 or pocket 2 from the best docking pose. These fragments exhibited extra interactions with CDK7 outside the core pocket that could enhance the binding affinity of novel inhibitors.
4. Second round of molecular docking. The extracted fragments were re-docked into pocket 1 or pocket 2 using smina to refine their conformations. We ranked them by smina scoring function, and retained the fragments with a smina affinity score within 1.2 kcal/mol from the best ligand.
5. Linker generation and screening. Linkers connecting fragments in the core pocket and pocket 1 or pocket 2 were generated using DiffLinker^45^. After manually removing candidates with a structurally invalid linker, 46 candidates remained for MD simulation (Table S2).
6. Stability checks with MD simulation. Each candidate molecule underwent a 200 ns MD simulations. To identify the most stable ligands, we calculated the standard deviation of root mean square deviation (RMSD) of the ligands over the final 50 ns of simulations and retained the 17 molecules with the lowest variations.

**Figure 2.**
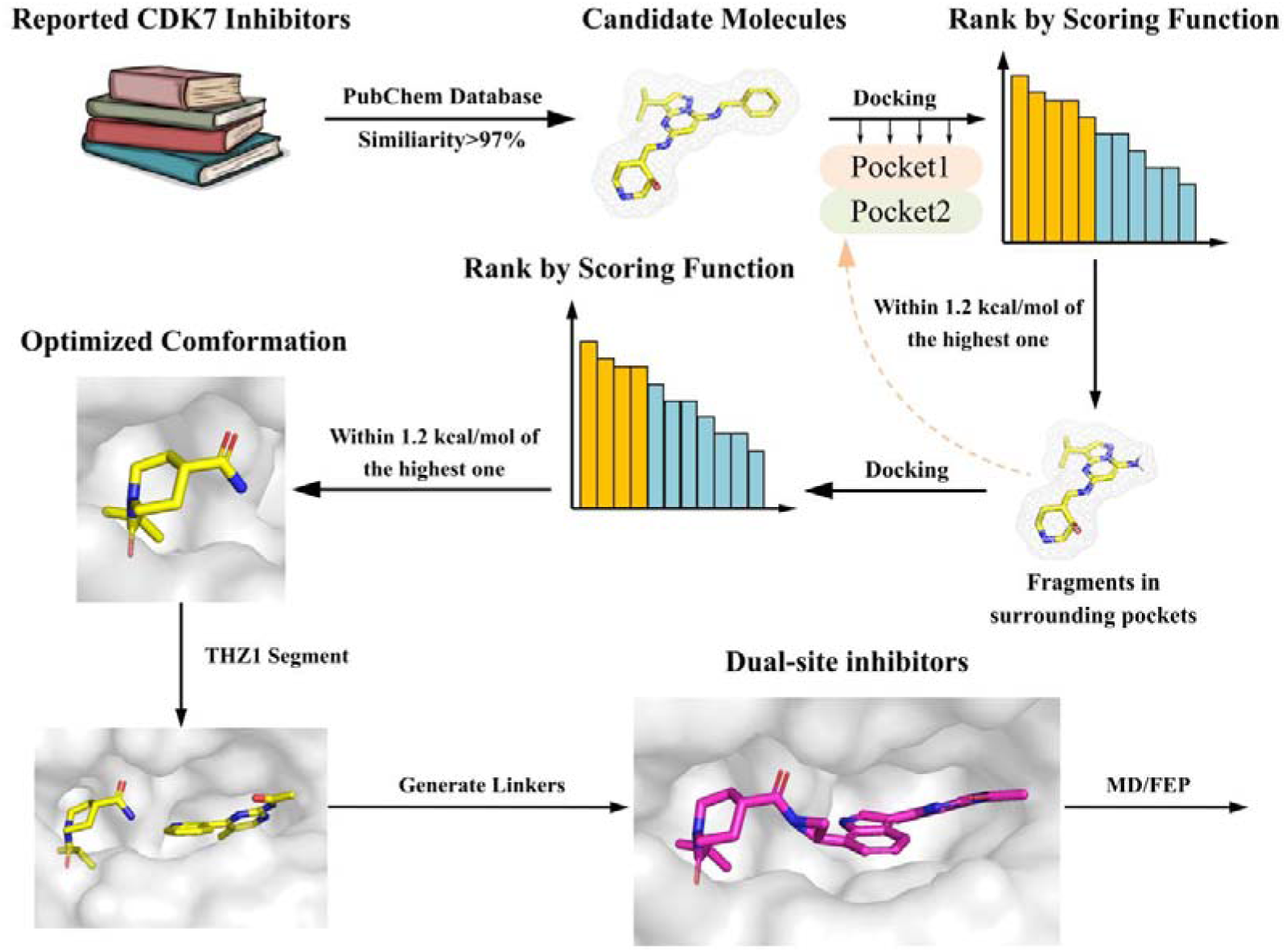
Virtual Screening Workflow for CDK7 Inhibitors.

The MMFF94 force field^46–50^ was applied to model small molecules during molecular docking.

### 2.3 All-Atom MD Simulation

The protein-ligand complexes were solvated with a 12 Å layer of TIP3P^51^ water, yielding a cubic box with approximately 31,000 atoms. To mimic physiological conditions, we added Na^+^ and Cl^-^ ions to reach a concentration of 0.15 M. All the system modeling was conducted in Visual Molecular Dynamics (VMD)^52^. The CAK complex, as well as Na^+^ and Cl^-^ ions were modeled using the CHARMM36 force field^53–56^. Parameters for all small molecules were generated in CHARMM-GUI^57–59^ using CHARMM General Force Field (CGenFF)^54^.

Energy minimization was performed in two stages using the default conjugate gradient algorithm: 5,000 steps with the protein backbone fixed, followed by 5,000 steps without constraints. The system was then heated to 310 K under an NVT ensemble in 90 ps, with position restraints of 1 kcal/(mol·Å²) applied to C-alpha atoms. Temperature was maintained at 310 K via Langevin dynamics with a damping coefficient of 1 ps^-^¹. The system was subsequently equilibrated under an NPT ensemble at 1.01325 bar using the Langevin piston method for 90 ps. Position restraints were progressively reduced (0.5, 0.25, 0.125, 0.0) over four 2-ns windows, followed by an additional 6 ns of unrestrained equilibration. We ran production simulations for 400 ns for the THZ1 derivative-CAK systems and 200 ns for candidate ligand-CAK systems.

All MD simulations were conducted with GPU-accelerated NAMD3^60^. Periodic boundary conditions were applied, with long-range electrostatics calculated using Particle Mesh Ewald (PME)^61^ with a grid spacing of 1 Å and a tolerance of 10^-6^. The van der Waals interactions were handled with a 12 Å cutoff, a switching distance of 10 Å, and a pair list cutoff of 13.5 Å. The SHAKE^62^ algorithm was used to constrain all bonds involving hydrogen atoms, enabling the use of a 2 fs integration time step. Additionally, we used the SETTLE^63^ algorithm to keep water molecules rigid. Coordinates were saved every 0.5 ns.

### 2.4 Trajectory Analysis

Trajectory analysis was conducted in VMD^52^. The RMSD of the ligand with the protein aligned was monitored to assess system stability throughout the simulation. Clustering analysis was conducted based on ligand RMSD with a 3.5 Å cutoff. Such a large cutoff ensures a single principal cluster even for the unstable THZ1-Noncov (Figure S2). The centroid of the largest cluster was extracted as the representative conformation for subsequent analysis and simulations. We used MDAnalysis^64,65^ to calculate the contact probability, ChimeraX^66–68^ and LigPlot^+^^69,70^ for protein-ligand interaction visualization.

### 2.5 Free Energy Perturbation (FEP) Simulations

FEP calculations were used to estimate the free energy difference (ΔΔG) between the designed molecules and the reference THZ1-Segment2. Two different thermodynamic cycle were used to compute the free energy difference between THZ1-Cov (Figure S1, right) or other molecules (Figure S1, left) and the reference molecule THZ1-Segment2. The theoretical framework for FEP is formulated as:

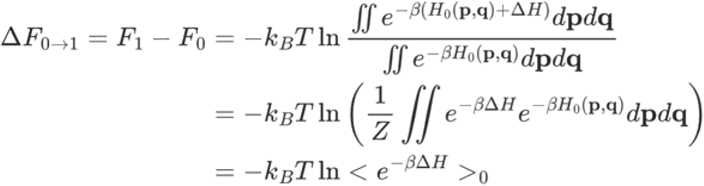

We prepared the dual topology files using our in-house software FEBuilder^71^. The soft-core potentials were incorporated to avoid endpoint artifacts. The coupling parameter (λ) varied across the following values: [0.00, 0.00001, 0.0001, 0.001, 0.01, 0.025, 0.05, 0.075, 0.1, 0.15, 0.2, 0.25, 0.3, 0.35, 0.4, 0.45, 0.5, 0.55, 0.6, 0.65, 0.7, 0.75, 0.8, 0.85, 0.9, 0.925, 0.95, 0.975, 0.99, 0.999, 0.9999, 0.99999, 1.00]. Each window underwent 5,000 steps of minimization, 20 ps of equilibration, followed by a 1 ns sampling. Other simulation parameters are the same as above. Each simulation was repeated 3 times to obtain the standard error of the free energy changes. The total time of the FEP calculations was 32 windows × 1 ns × 2 × 3 runs × 17 ligands = 3360 ns.

The workflow begins with a similarity search in the PubChem database, identifying 147 molecules with ≥97% similarity to eight known CDK7 inhibitors. These candidates were docked into CDK7’s binding site using smina. Fragments with favorable interactions in pocket 1 or pocket 2 were extracted and re-docked to refine conformations. High-scoring fragments were connected to THZ1-Segment2 with the DiffLinker model. They were checked manually to yield 46 final candidates for evaluation in MD and FEP simulations.

## 3 Results and Discussion

### 3.1 Structural basis of THZ1 binding to CAK

Using the structures of THZ1-Cov and THZ1-Noncov as starting points, we conducted 400 ns molecular dynamics (MD) simulations. RMSD analysis of the trajectories revealed that THZ1-Cov is significantly more stable than THZ1-Noncov. Specifically, THZ1-Cov maintained stable binding to the CAK complex, with minimal conformational changes from the crystal structure (Figure 3B). The covalent bond constrained the flexible regions of THZ1 outside the core pocket, stabilizing interactions with residues such as Cys312 and Asn311 (Figure 4). In contrast, THZ1-Noncov exhibited pronounced conformational fluctuations, particularly in regions outside the core pocket (Figure 3C). The non-covalent form interacted less frequently with Cys312 and more frequently with Phe17, an interaction not observed in THZ1-Cov (Figure 4). This reduced stability likely explains the weaker binding efficacy of THZ1-Noncov to CDK7 (146 nM) compared to THZ1-Cov (32 nM)^72^ in experiments.

**Figure 3.**
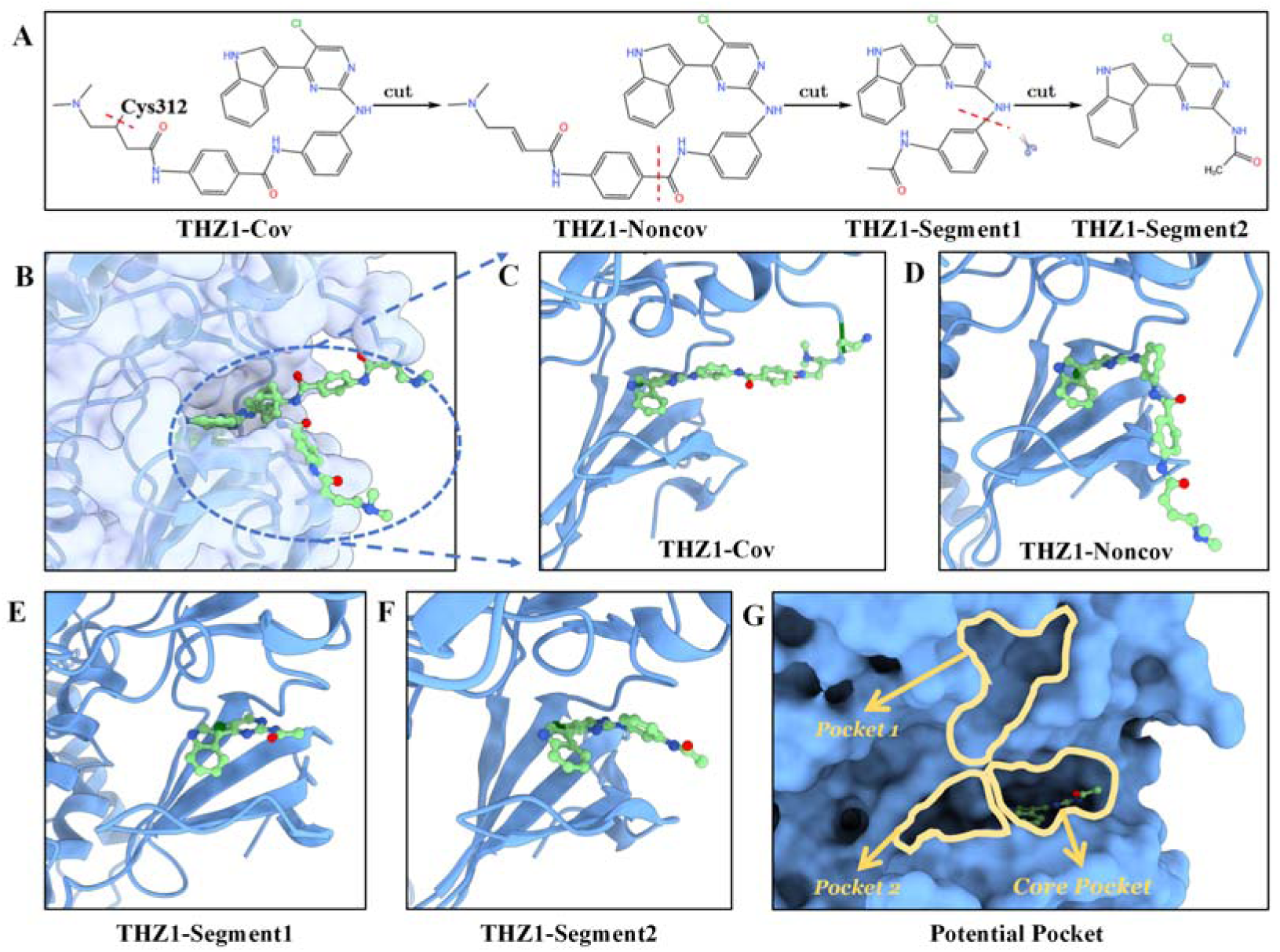
Structural analysis of THZ1-CAK complex. CDK7 is shown as a blue cartoon, while small molecules are represented as green ball-and-stick models. (A) Schematic depiction of THZ1 divided into covalent (THZ1-Cov) and non-covalent (THZ1-Noncov) forms, and two simplified segments: THZ1-Segment1 and THZ1-Segment2. Amino groups are capped with acetyl group. (B) Overlay of representative conformations of THZ1-Cov and THZ1-Noncov within the CDK7 binding pocket from clustering analysis of MD simulations. THZ1-Cov forms a stable covalent bond with Cys312, while THZ1-Noncov exhibits flexibility outside the core pocket. (C-D) Detailed views of THZ1-Cov and THZ1-Noncov in the CDK7 pocket, demonstrating distinct binding conformations. (E-F) Representative conformations of THZ1-Segment1 and THZ1-Segment2, highlighting their respective stabilities within the CDK7 pocket. (G) Binding pockets around CDK7 binding site indicating potential binding pockets for future screening. The core and surrounding pockets are outlined in yellow and orange, respectively.

**Figure 4.**
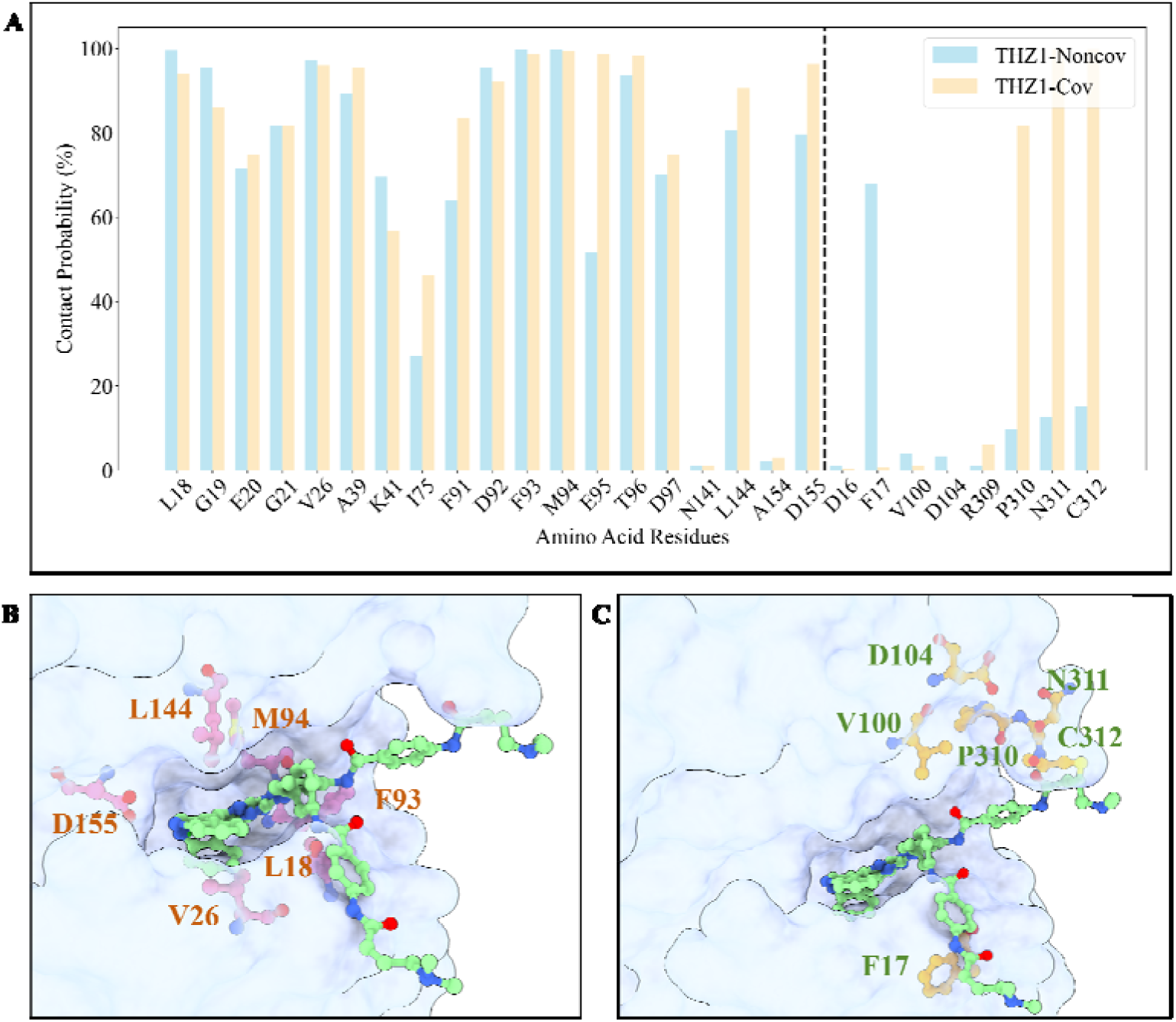
Contact analysis of THZ1 with CDK7. CDK7 is shown as a light blue surface, with small molecules depicted as green ball-and-stick models. (A) Contact probability between THZ1-Cov (blue) and THZ1-Noncov (orange) with CDK7 residues, categorized as the core pocket (left of the dashed line) and the covalent region (right of the dashed line). (B) Both molecules form frequent contacts with core residues Leu18, Val26, Phe93, Leu144, and Asp155. Asp155 forms a hydrogen bond with the indole NH group of THZ1, while other residues primarily contribute to hydrophobic interactions. (C) THZ1-Cov is stabilized by the covalent bond with Cys312. THZ1-Noncov may display interactions with distant residues like Phe17, indicating greater conformational flexibility.

Despite higher RMSD fluctuations, THZ1-Noncov retained interactions within the core pocket similar to those of THZ1-Cov. In particular, it formed frequent and stable contacts with residues Leu18, Val26, Phe93, Leu144, and Asp155. Notably, the carboxyl group in Asp155 formed a hydrogen bond with the indole NH group, while the backbone of Met94 formed a hydrogen bond with the linker NH group. Additional interactions were predominantly hydrophobic (Figure 4B). These interactions underscore the importance of the core pocket in stabilizing THZ1 binding. The stability of THZ1-Noncov within the core pocket likely accounts for its residual binding to CDK7 despite the absence of a covalent bond.

To further investigate these interactions, we took two fragments of THZ1 —THZ1-Segment1 and THZ1-Segment2 (Figure 3A)— and subjected each of them to 400 ns MD simulations within the CAK complex (Figures 3E-F). Both fragments, particularly THZ1-Segment2, exhibited stable binding with minimal conformational changes (Figure S2), closely resembling the binding patterns of THZ1-Cov and THZ1-Noncov. These results highlight the crucial role of the core pocket in CDK7 binding and suggest potential avenues for inhibitor design.

THZ1-Segment2 emerges as a promising scaffold for development of CDK7 inhibitors. By replacing the covalent region with non-covalent interactions that provide comparable binding affinity, it may be possible to design novel and selective CDK7 inhibitors.

### 3.2 Enhancing CDK7 Binding by Integrating Fragments from Known Inhibitors

To enhance non-covalent interactions for CDK7 inhibition, we analyzed key molecule fragments derived from eight known CDK7 inhibitors—LDC4297, SY5609, Roscovitine, CT7001, YKL-5-124, SNS-032, SY-1365, and Flavopiridol. A similarity search in the PubChem database yielded 147 molecules, including the known inhibitors. These molecules were docked into a box encompassing the core pocket and adjacent pockets (pocket 1 and pocket 2; Figure 3G). Binding affinities, represented as smina scores (in kcal/mol), were used to rank the molecules, with lower scores indicating stronger binding. Molecules with smina scores within 1.2 kcal/mol of the best ligand (i.e., below −9.5 kcal/mol; Figure S3) were selected for further analysis. From this subset, molecular fragments binding to pocket 1 or pocket 2 were extracted, with most fragments originating from YKL-5-124 and SY-1365.

The extracted fragments were re-docked to refine their conformations and improve binding affinity. We obtained fragments with a smina score difference of less than 1.2 kcal/mol from the best pose and a relatively small deviation from their initial pose. To construct the complete ligands, the deep learning program DiffLinker was used to generate linkers between the selected fragments and THZ1-Segment2, taking pocket geometry into account. After manual inspection to eliminate distorted structures, we identified 46 candidates as potential CDK7 inhibitors (CDI; Table S1).

To assess the structural stability of these candidates, we performed 200 ns MD simulations to relax their conformations. Candidates were ranked based on the standard deviation of their RMSD values over the final 50 ns of the simulations (Figure S4). The top 40% of candidates, comprising 17 ligands, exhibited minimal conformational changes and were advanced to free energy perturbation (FEP) calculations as potential CDK7 inhibitors. Among these, CDI7, CDI8, CDI9, CDI18, and CDI29 predominantly occupied pocket 1, while CDI1, CDI3, CDI5, CDI21, CDI27, CDI32, CDI33, CDI36, CDI37, CDI42, CDI48, and CDI50 occupied pocket 2. The preference for pocket 2 as a binding site is likely attributed to its deeper cavity, which provides additional stabilization.

### 3.3 Identification of Potential CDK7 Inhibitors through FEP Calculations

Since THZ1 forms a covalent bond with Cys312 in the CAK complex, it was necessary to calculate the free energy change associate with this bond formation to compare the binding free energies of covalent and non-covalent ligands. The thermodynamics cycles used for these calculations are shown in Figure S1. Based on density functional theory (DFT) calculations by Krenske et al., the free energy change for a cysteine residue covalently binding to a molecule similar to THZ1 (referred to as molecule 18 in their work) via Michael addition was estimated at −5.7 kcal/mol (Figure S5)^73^. While THZ1 differs from molecule 18 in that its trialkylammonium cation is replaced by a cyclopropyl group and its amide by a methyl ester, these structural differences are unlikely to significantly alter the reaction energetics. Although the cyclopropyl group may slightly reduce reaction efficiency, this approximation provides a reasonable estimate for the free energy change transforming THZ1-Cov into THZ1-Noncov.

Since THZ1-Noncov and all candidate molecules share the THZ1-Segment2 fragment, we computed their binding free energies relative to THZ1-Segment2. In particular, FEP calculations were performed to transform each designed molecule into THZ1-Segment2, with higher ΔΔG values indicating stronger binding. Our results revealed that most fragments identified through virtual screening possess an enhanced binding affinity (Figure 5A). Using the ΔΔG value for the transformation of THZ1-Cov to THZ1-Segment2 (Figure S1) as a baseline, we compared all candidate molecules to THZ1-Cov. Notably, CDI3, CDI18, and CDI37 displayed significantly higher ΔΔG values relative to the baseline while maintaining stable binding conformations (Figure 5B). Other candidates, such as CDI7 and CDI29, demonstrated ΔΔG values comparable to the baseline, suggesting potential efficacy. The error bars in Figure 5A represent standard deviations, reflecting inherent variability in FEP results due to conformational sampling. Despite this variability, the consistently high ΔΔG values of CDI18 and CDI37 suggest robust binding compare to the baseline.

**Figure 5.**
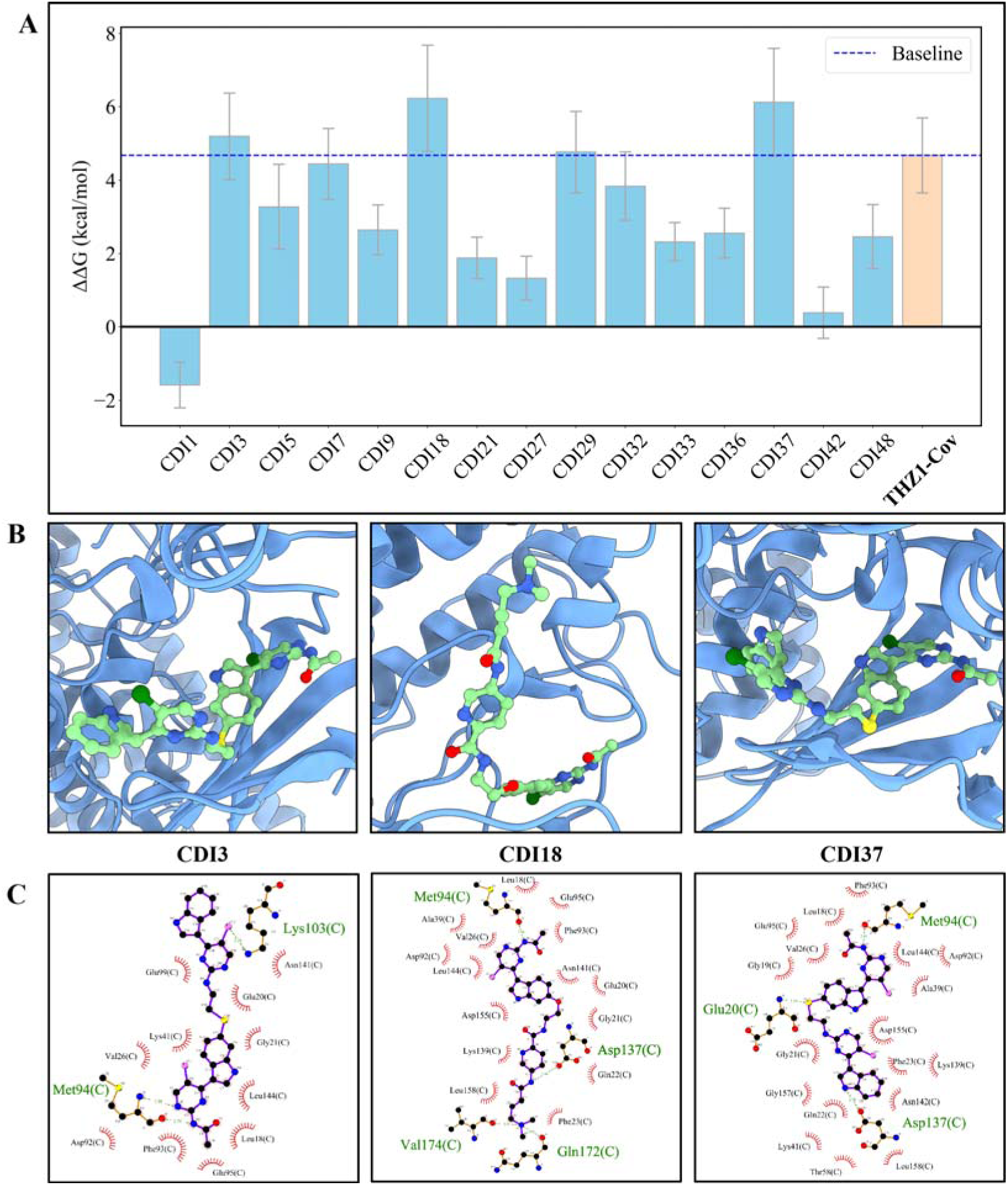
Relative binding free energy and interaction profiles of designed CDK7 inhibitors. CDK7 is illustrated in a blue cartoon representation, and the inhibitors are depicted as green ball-and-stick models. (A) Bar plot of ΔΔG for various CDIs (blue) transforming into THZ1-Segment2. Error bars indicate standard deviations. THZ1-Cov (orange) is the baseline, accounting for covalent bond formation energy plus ΔΔG transforming THZ1-Noncov to THZ1-Segment2. Positive ΔΔG values indicate higher binding affinity, with CDI3, CDI18, and CDI37 exhibiting the strongest binding affinities. (B) Representative binding conformations of the top candidates—CDI3, CDI18, and CDI37—within the CDK7 binding pocket. The THZ1-Segment2 fragments in them maintain a stable conformation in the core pocket across different inhibitors. These conformations are derived from clustering analysis to serve as the starting conformations for FEP calculations. (C) Detailed two-dimensional interaction maps for CDI3, CDI18, and CDI37 generated by LigPlot^+^, illustrating extensive non-polar interactions, hydrogen bonding, and salt bridges that contribute to enhanced binding stability.

Figure 5C illustrates the detailed interactions of the top candidate inhibitors within the core pocket and pocket 2. In the core pocket, THZ1-Segment2 forms a stable hydrogen bond with the carbonyl group of Met94, serving as an anchor. Additional hydrophobic interactions with Leu18, Val26, Phe93, and Leu144, along with polar contacts with Asp92 and Asp155, further enhance binding stability. These interactions closely resemble those observed in Figure 4A. Furthermore, linker regions establish favorable contacts with Glu20 and Gly21, further contributing to overall binding stability.

CDI3 and CDI37 exhibit structural similarity with subtle differences. CDI3 includes a double-bonded nitrogen in the linker and a saturated five-membered ring in pocket 2, forming key interactions with residues Glu99, Lys103, and Asn141. CDI37, which feature a saturated linker and a purine group, engages with a broader network of interactions, including contacts with Gln22, Phe93, and Asn142. Its purine group forms a particularly stable hydrogen bond with Asp137, significantly enhancing binding within pocket 2. CDI18, which structurally resembles THZ1 but with a shorter linker, primarily inserts into pocket 1. Key interaction in pocket 1 involve hydrogen bonding between its amide group and Asp137, as well as interactions with Lys139 and Leu158. Additionally, its trialkylammonium group interacts with the backbone carbonyl groups of Gln172 and Val174, and the side chain of Phe23, through electrostatic and π-cation interactions, further strengthening its binding.

In summary, the robust networks of polar and non-polar interactions observed in CDI3, CDI18, and CDI37 support their potential as potent CDK7 inhibitors, making them promising candidates for further development.

## 4. Conclusion

In this study, we established a comprehensive workflow that integrates virtual screening, molecular dynamics simulations, and free energy perturbation calculations to identify potential non-covalent small molecule inhibitors targeting CDK7. Our findings highlight the pivotal role of the THZ1-Segment2 fragment, which demonstrated stable interactions within the CDK7 ATP-binding pocket, positioning it as a promising scaffold for novel inhibitor design. While the covalent bond of THZ1 plays a crucial role in binding stability, our innovative focus on non-covalent compounds offers distinct advantages, including reduced off-target effects and improved selectivity. By incorporating molecular fragments from known inhibitors, we identified three high-affinity compounds, providing valuable insights for future development of CDK inhibitors. Moreover, our overall workflow can be further automated, thereby facilitating efficient drug discovery for other targets.

Despite the effectiveness of our computational simulation methodology, we acknowledge several inherent limitations. First, the fragment library used in this study was limited to known inhibitors assumed to bind in pocket 1 and/or pocket 2, which may constrain the diversity of novel fragments interactions. Expanding the library to include novel chemical structures will enhance interaction diversity and improve hit identification. Second, our approximation of covalent binding energetics may not fully account for the influence of substituent groups in the Michael addition receptor. Future work will involve more rigorous quantum mechanical calculations to better model these effects. These planned refinements will enhance the robustness and confidence of our workflow, strengthening its utility for inhibitor design.

The computational nature of this study also emphasizes the necessity of experimental validation to confirm the binding affinities and biological efficacy of the identified compounds. For example, it is highly desired to have experiments focusing on binding affinity measurements via surface plasmon resonance (SPR), cytotoxicity evaluations in cell-based assays, and comprehensive anti-tumor efficacy tests in mouse models. Insights from these studies will guide the optimization of pharmacological properties, reduce potential side effects, and refine the inhibitors’ overall therapeutic potential. Our current work provides the foundation for advancement in future development of CDK7-targeted therapeutics, contributing to the discovery of more effecting and selective treatments for various cancers.

## Supporting information

Cover Letter

## Acknowledgements

We thank Liquan Huang, Lianxue Zhang, Yanqing Yang, and Yangwei Jiang for helpful discussions. This work was partially supported by the National Key R&D Program of China (2021YFF1200404 and 2021YFA1201200), the National Natural Science Foundation of China (U1967217), the National Center of Technology Innovation for Biopharmaceuticals (NCTIB2022HS02010), Shanghai Artificial Intelligence Lab (P22KN00272), the National Independent Innovation Demonstration Zone Shanghai Zhangjiang Major Projects (ZJZX2020014), the Starry Night Science Fund of Zhejiang University Shanghai Institute for Advanced Study (SN-ZJU-SIAS-003) and Zhejiang University Global Partnership Fund (188170+194452409/004).

## References

(1) Sava, G. P.; Fan, H.; Coombes, R. C.; Buluwela, L.; Ali, S. CDK7 Inhibitors as Anticancer Drugs. Cancer Metastasis Rev. 2020, 39 (3), 805–823. 10.1007/s10555-020-09885-8.

(2) Akoulitchev, S.; Mäkelä, T. P.; Weinberg, R. A.; Reinberg, D. Requirement for TFIIH Kinase Activity in Transcription by RNA Polymerase II. Nature 1995, 377 (6549), 557–560. 10.1038/377557a0.

(3) Ko, L. J.; Shieh, S.-Y.; Chen, X.; Jayaraman, L.; Tamai, K.; Taya, Y.; Prives, C.; Pan, Z.-Q. P53 Is Phosphorylated by CDK7-Cyclin H in a P36 *^MAT1^* -Dependent Manner. Mol. Cell. Biol. 1997, 17 (12), 7220–7229. 10.1128/MCB.17.12.7220.

(4) Chen, J.; Larochelle, S.; Li, X.; Suter, B. Xpd/Ercc2 Regulates CAK Activity and Mitotic Progression. Nature 2003, 424 (6945), 228–232. 10.1038/nature01746.

(5) Malumbres, M. Cyclin-Dependent Kinases. Genome Biol. 2014, 15 (6), 122. 10.1186/gb4184.

(6) Schachter, M. M.; Merrick, K. A.; Larochelle, S.; Hirschi, A.; Zhang, C.; Shokat, K. M.; Rubin, S. M.; Fisher, R. P. A Cdk7-Cdk4 T-Loop Phosphorylation Cascade Promotes G1 Progression. Mol. Cell 2013, 50 (2), 250–260. 10.1016/j.molcel.2013.04.003.

(7) Larochelle, S.; Merrick, K. A.; Terret, M.-E.; Wohlbold, L.; Barboza, N. M.; Zhang, C.; Shokat, K. M.; Jallepalli, P. V.; Fisher, R. P. Requirements for Cdk7 in the Assembly of Cdk1/Cyclin B and Activation of Cdk2 Revealed by Chemical Genetics in Human Cells. Mol. Cell 2007, 25 (6), 839–850. 10.1016/j.molcel.2007.02.003.

(8) Lapenna, S.; Giordano, A. Cell Cycle Kinases as Therapeutic Targets for Cancer. Nat. Rev. Drug Discov. 2009, 8 (7), 547–566. 10.1038/nrd2907.

(9) Gong, Y.; Li, H. CDK7 in Breast Cancer: Mechanisms of Action and Therapeutic Potential. Cell Commun. Signal. 2024, 22 (1), 226. 10.1186/s12964-024-01577-y.

(10) Wang, Y.; Zhang, T.; Kwiatkowski, N.; Abraham, B. J.; Lee, T. I.; Xie, S.; Yuzugullu, H.; Von, T.; Li, H.; Lin, Z.; Stover, D. G.; Lim, E.; Wang, Z. C.; Iglehart, J. D.; Young, R. A.; Gray, N. S.; Zhao, J. J. CDK7-Dependent Transcriptional Addiction in Triple-Negative Breast Cancer. Cell 2015, 163 (1), 174–186. 10.1016/j.cell.2015.08.063.

(11) Hur, J. Y.; Kim, H. R.; Lee, J. Y.; Park, S.; Hwang, J. A.; Kim, W. S.; Yoon, S.; Choi, C.-M.; Rho, J. K.; Lee, J. C. CDK7 Inhibition as a Promising Therapeutic Strategy for Lung Squamous Cell Carcinomas with a SOX2 Amplification. Cell. Oncol. 2019, 42 (4), 449–458. 10.1007/s13402-019-00434-2.

(12) Cao, X.; Dang, L.; Zheng, X.; Lu, Y.; Lu, Y.; Ji, R.; Zhang, T.; Ruan, X.; Zhi, J.; Hou, X.; Yi, X.; Li, M. J.; Gu, T.; Gao, M.; Zhang, L.; Chen, Y. Targeting Super-Enhancer-Driven Oncogenic Transcription by CDK7 Inhibition in Anaplastic Thyroid Carcinoma. Thyroid 2019, 29 (6), 809–823. 10.1089/thy.2018.0550.

(13) Wang, M.; Wang, T.; Zhang, X.; Wu, X.; Jiang, S. Cyclin-Dependent Kinase 7 Inhibitors in Cancer Therapy. Future Med. Chem. 2020, 12 (9), 813–833. 10.4155/fmc-2019-0334.

(14) Li, Z.-M.; Liu, G.; Gao, Y.; Zhao, M.-G. Targeting CDK7 in Oncology: The Avenue Forward. Pharmacol. Ther. 2022, 240, 108229. 10.1016/j.pharmthera.2022.108229.

(15) Sedlacek, H.; Czech, J.; Naik, R.; Kaur, G.; Worland, P.; Losiewicz, M.; Parker, B.; Carlson, B.; Smith, A.; Senderowicz, A.; Sausville, E. Flavopiridol (L86 8275; NSC 649890), a New Kinase Inhibitor for Tumor Therapy. Int. J. Oncol. 1996. 10.3892/ijo.9.6.1143.

(16) Chen, R.; Wierda, W. G.; Chubb, S.; Hawtin, R. E.; Fox, J. A.; Keating, M. J.; Gandhi, V.; Plunkett, W. Mechanism of Action of SNS-032, a Novel Cyclin-Dependent Kinase Inhibitor, in Chronic Lymphocytic Leukemia. Blood 2009, 113 (19), 4637–4645. 10.1182/blood-2008-12-190256.

(17) Meijer, L.; Raymond, E. Roscovitine and Other Purines as Kinase Inhibitors. From Starfish Oocytes to Clinical Trials. Acc. Chem. Res. 2003, 36 (6), 417–425. 10.1021/ar0201198.

(18) Ali, S.; Heathcote, D. A.; Kroll, S. H. B.; Jogalekar, A. S.; Scheiper, B.; Patel, H.; Brackow, J.; Siwicka, A.; Fuchter, M. J.; Periyasamy, M.; Tolhurst, R. S.; Kanneganti, S. K.; Snyder, J. P.; Liotta, D. C.; Aboagye, E. O.; Barrett, A. G. M.; Coombes, R. C. The Development of a Selective Cyclin-Dependent Kinase Inhibitor That Shows Antitumor Activity. Cancer Res. 2009, 69 (15), 6208–6215. 10.1158/0008-5472.CAN-09-0301.

(19) Peng, J.; Yang, M.; Bi, R.; Wang, Y.; Wang, C.; Wei, X.; Zhang, Z.; Xie, X.; Wei, W. Targeting Mutated P53 Dependency in Triple-Negative Breast Cancer Cells Through CDK7 Inhibition. Front. Oncol. 2021, 11, 664848. 10.3389/fonc.2021.664848.

(20) Zhang, H.; Christensen, C. L.; Dries, R.; Oser, M. G.; Deng, J.; Diskin, B.; Li, F.; Pan, Y.; Zhang, X.; Yin, Y.; Papadopoulos, E.; Pyon, V.; Thakurdin, C.; Kwiatkowski, N.; Jani, K.; Rabin, A. R.; Castro, D. M.; Chen, T.; Silver, H.; Huang, Q.; Bulatovic, M.; Dowling, C. M.; Sundberg, B.; Leggett, A.; Ranieri, M.; Han, H.; Li, S.; Yang, A.; Labbe, K. E.; Almonte, C.; Sviderskiy, V. O.; Quinn, M.; Donaghue, J.; Wang, E. S.; Zhang, T.; He, Z.; Velcheti, V.; Hammerman, P. S.; Freeman, G. J.; Bonneau, R.; Kaelin, W. G.; Sutherland, K. D.; Kersbergen, A.; Aguirre, A. J.; Yuan, G.-C.; Rothenberg, E.; Miller, G.; Gray, N. S.; Wong, K.-K. CDK7 Inhibition Potentiates Genome Instability Triggering Anti-Tumor Immunity in Small Cell Lung Cancer. Cancer Cell 2020, 37 (1), 37–54.e9. 10.1016/j.ccell.2019.11.003.

(21) Panagiotou, E.; Gomatou, G.; Trontzas, I. P.; Syrigos, N.; Kotteas, E. Cyclin-Dependent Kinase (CDK) Inhibitors in Solid Tumors: A Review of Clinical Trials. Clin. Transl. Oncol. 2022, 24 (2), 161–192. 10.1007/s12094-021-02688-5.

(22) Zheng, L.; Li, Y.; Wu, D.; Xiao, H.; Zheng, S.; Wang, G.; Sun, Q. Development of Covalent Inhibitors: Principle, Design, and Application in Cancer. MedComm – Oncol. 2023, 2 (4), e56. 10.1002/mog2.56.

(23) Kwiatkowski, N.; Zhang, T.; Rahl, P. B.; Abraham, B. J.; Reddy, J.; Ficarro, S. B.; Dastur, A.; Amzallag, A.; Ramaswamy, S.; Tesar, B.; Jenkins, C. E.; Hannett, N. M.; McMillin, D.; Sanda, T.; Sim, T.; Kim, N. D.; Look, T.; Mitsiades, C. S.; Weng, A. P.; Brown, J. R.; Benes, C. H.; Marto, J. A.; Young, R. A.; Gray, N. S. Targeting Transcription Regulation in Cancer with a Covalent CDK7 Inhibitor. Nature 2014, 511 (7511), 616–620. 10.1038/nature13393.

(24) Kwiatkowski, N.; Zhang, T.; Rahl, P. B.; Abraham, B. J.; Reddy, J.; Ficarro, S. B.; Dastur, A.; Amzallag, A.; Ramaswamy, S.; Tesar, B.; Jenkins, C. E.; Hannett, N. M.; McMillin, D.; Sanda, T.; Sim, T.; Kim, N. D.; Look, T.; Mitsiades, C. S.; Weng, A. P.; Brown, J. R.; Benes, C. H.; Marto, J. A.; Young, R. A.; Gray, N. S. Targeting Transcription Regulation in Cancer with a Covalent CDK7 Inhibitor. Nature 2014, 511 (7511), 616–620. 10.1038/nature13393.

(25) Spagnolo, P.; Kropski, J. A.; Jones, M. G.; Lee, J. S.; Rossi, G.; Karampitsakos, T.; Maher, T. M.; Tzouvelekis, A.; Ryerson, C. J. Idiopathic Pulmonary Fibrosis: Disease Mechanisms and Drug Development. Pharmacol. Ther. 2021, 222, 107798. 10.1016/j.pharmthera.2020.107798.

(26) Kuo, K.-L.; Lin, W.-C.; Liu, S.-H.; Hsu, F.-S.; Kuo, Y.; Liao, S.-M.; Yang, S.-P.; Wang, Z.-H.; Hsu, C.-H.; Huang, K.-H. THZ1, a Covalent CDK7 Inhibitor, Enhances Gemcitabine-Induced Cytotoxicity via Suppression of Bcl-2 in Urothelial Carcinoma. Am. J. Cancer Res. 2021, 11 (1), 171–180.

(27) Zhang, J.; Liu, W.; Zou, C.; Zhao, Z.; Lai, Y.; Shi, Z.; Xie, X.; Huang, G.; Wang, Y.; Zhang, X.; Fan, Z.; Su, Q.; Yin, J.; Shen, J. Targeting Super-Enhancer-Associated Oncogenes in Osteosarcoma with THZ2, a Covalent CDK7 Inhibitor. Clin. Cancer Res. Off. J. Am. Assoc. Cancer Res. 2020, 26 (11), 2681–2692. 10.1158/1078-0432.CCR-19-1418.

(28) Sharifnia, T.; Wawer, M. J.; Chen, T.; Huang, Q.-Y.; Weir, B. A.; Sizemore, A.; Lawlor, M. A.; Goodale, A.; Cowley, G. S.; Vazquez, F.; Ott, C. J.; Francis, J. M.; Sassi, S.; Cogswell, P.; Sheppard, H. E.; Zhang, T.; Gray, N. S.; Clarke, P. A.; Blagg, J.; Workman, P.; Sommer, J.; Hornicek, F.; Root, D. E.; Hahn, W. C.; Bradner, J. E.; Wong, K. K.; Clemons, P. A.; Lin, C. Y.; Kotz, J. D.; Schreiber, S. L. Small-Molecule Targeting of Brachyury Transcription Factor Addiction in Chordoma. Nat. Med. 2019, 25 (2), 292–300. 10.1038/s41591-018-0312-3.

(29) Christensen, C. L.; Kwiatkowski, N.; Abraham, B. J.; Carretero, J.; Al-Shahrour, F.; Zhang, T.; Chipumuro, E.; Herter-Sprie, G. S.; Akbay, E. A.; Altabef, A.; Zhang, J.; Shimamura, T.; Capelletti, M.; Reibel, J. B.; Cavanaugh, J. D.; Gao, P.; Liu, Y.; Michaelsen, S. R.; Poulsen, H. S.; Aref, A. R.; Barbie, D. A.; Bradner, J. E.; George, R. E.; Gray, N. S.; Young, R. A.; Wong, K.-K. Targeting Transcriptional Addictions in Small Cell Lung Cancer with a Covalent CDK7 Inhibitor. Cancer Cell 2014, 26 (6), 909–922. 10.1016/j.ccell.2014.10.019.

(30) Gao, Y.; Zhang, T.; Terai, H.; Ficarro, S. B.; Kwiatkowski, N.; Hao, M.-F.; Sharma, B.; Christensen, C. L.; Chipumuro, E.; Wong, K.-K.; Marto, J. A.; Hammerman, P. S.; Gray, N. S.; George, R. E. Overcoming Resistance to the THZ Series of Covalent Transcriptional CDK Inhibitors. Cell Chem. Biol. 2018, 25 (2), 135–142.e5. 10.1016/j.chembiol.2017.11.007.

(31) Chen, H.-R.; Lin, G.-T.; Huang, C.-K.; Fann, M.-J. Cdk12 and Cdk13 Regulate Axonal Elongation through a Common Signaling Pathway That Modulates Cdk5 Expression. Exp. Neurol. 2014, 261, 10–21. 10.1016/j.expneurol.2014.06.024.

(32) Echalier, A.; Hole, A. J.; Lolli, G.; Endicott, J. A.; Noble, M. E. M. An Inhibitor’s-Eye View of the ATP-Binding Site of CDKs in Different Regulatory States. ACS Chem. Biol. 2014, 9 (6), 1251–1256. 10.1021/cb500135f.

(33) Olson, C. M.; Liang, Y.; Leggett, A.; Park, W. D.; Li, L.; Mills, C. E.; Elsarrag, S. Z.; Ficarro, S. B.; Zhang, T.; Düster, R.; Geyer, M.; Sim, T.; Marto, J. A.; Sorger, P. K.; Westover, K. D.; Lin, C. Y.; Kwiatkowski, N.; Gray, N. S. Development of a Selective CDK7 Covalent Inhibitor Reveals Predominant Cell-Cycle Phenotype. Cell Chem. Biol. 2019, 26 (6), 792–803.e10. 10.1016/j.chembiol.2019.02.012.

(34) Diab, S.; Yu, M.; Wang, S. CDK7 Inhibitors in Cancer Therapy: The Sweet Smell of Success? J. Med. Chem. 2020, 63 (14), 7458–7474. 10.1021/acs.jmedchem.9b01985.

(35) Düster, R.; Anand, K.; Binder, S. C.; Schmitz, M.; Gatterdam, K.; Fisher, R. P.; Geyer, M. Structural Basis of Cdk7 Activation by Dual T-Loop Phosphorylation. Nat. Commun. 2024, 15 (1), 6597. 10.1038/s41467-024-50891-z.

(36) Song, Y.; Bell, D. R.; Ahmed, R.; Chan, K. C.; Lee, S.; Hamad, A. R. A.; Zhou, R. A Mutagenesis Study of Autoantigen Optimization for Potential T1D Vaccine Design. Proc. Natl. Acad. Sci. 2023, 120 (16), e2214430120. 10.1073/pnas.2214430120.

(37) Chowell, D.; Morris, L. G. T.; Grigg, C. M.; Weber, J. K.; Samstein, R. M.; Makarov, V.; Kuo, F.; Kendall, S. M.; Requena, D.; Riaz, N.; Greenbaum, B.; Carroll, J.; Garon, E.; Hyman, D. M.; Zehir, A.; Solit, D.; Berger, M.; Zhou, R.; Rizvi, N. A.; Chan, T. A. Patient HLA Class I Genotype Influences Cancer Response to Checkpoint Blockade Immunotherapy. Science 2018, 359 (6375), 582–587. 10.1126/science.aao4572.

(38) Ahmed, R.; Omidian, Z.; Giwa, A.; Cornwell, B.; Majety, N.; Bell, D. R.; Lee, S.; Zhang, H.; Michels, A.; Desiderio, S.; Sadegh-Nasseri, S.; Rabb, H.; Gritsch, S.; Suva, M. L.; Cahan, P.; Zhou, R.; Jie, C.; Donner, T.; Hamad, A. R. A. A Public BCR Present in a Unique Dual-Receptor-Expressing Lymphocyte from Type 1 Diabetes Patients Encodes a Potent T Cell Autoantigen. Cell 2019, 177 (6), 1583–1599.e16. 10.1016/j.cell.2019.05.007.

(39) Bieniossek, C.; Richmond, T. J.; Berger, I. MultiBac: Multigene BaculovirusCBased Eukaryotic Protein Complex Production. Curr. Protoc. Protein Sci. 2008, 51 (1). 10.1002/0471140864.ps0520s51.

(40) Glover-Cutter, K.; Larochelle, S.; Erickson, B.; Zhang, C.; Shokat, K.; Fisher, R. P.; Bentley, D. L. TFIIH-Associated Cdk7 Kinase Functions in Phosphorylation of C-Terminal Domain Ser7 Residues, Promoter-Proximal Pausing, and Termination by RNA Polymerase II. Mol. Cell. Biol. 2009, 29 (20), 5455–5464. 10.1128/MCB.00637-09.

(41) Waterhouse, A.; Bertoni, M.; Bienert, S.; Studer, G.; Tauriello, G.; Gumienny, R.; Heer, F. T.; de Beer, T. A. P.; Rempfer, C.; Bordoli, L.; Lepore, R.; Schwede, T. SWISS-MODEL: Homology Modelling of Protein Structures and Complexes. Nucleic Acids Res. 2018, 46 (W1), W296–W303. 10.1093/nar/gky427.

(42) Kim, S.; Chen, J.; Cheng, T.; Gindulyte, A.; He, J.; He, S.; Li, Q.; Shoemaker, B. A.; Thiessen, P. A.; Yu, B.; Zaslavsky, L.; Zhang, J.; Bolton, E. E. PubChem 2023 Update. Nucleic Acids Res. 2023, 51 (D1), D1373–D1380. 10.1093/nar/gkac956.

(43) Koes, D. R.; Baumgartner, M. P.; Camacho, C. J. Lessons Learned in Empirical Scoring with Smina from the CSAR 2011 Benchmarking Exercise. J. Chem. Inf. Model. 2013, 53 (8), 1893–1904. 10.1021/ci300604z.

(44) Chern, C. J.; Beutler, E. Biochemical and Electrophoretic Studies of Erythrocyte Pyridoxine Kinase in White and Black Americans. Am. J. Hum. Genet. 1976, 28 (1), 9–17.

(45) Igashov, I.; Stärk, H.; Vignac, C.; Schneuing, A.; Satorras, V. G.; Frossard, P.; Welling, M.; Bronstein, M.; Correia, B. Equivariant 3D-Conditional Diffusion Model for Molecular Linker Design. *Nat*. Mach. Intell. 2024, 6 (4), 417–427. 10.1038/s42256-024-00815-9.

(46) Halgren, T. A. Merck Molecular Force Field. I. Basis, Form, Scope, Parameterization, and Performance of MMFF94. J. Comput. Chem. 1996, 17 (5–6), 490–519. 10.1002/(SICI)1096-987X(199604)17:5/6<490::AID-JCC1>3.0.CO;2-P.

(47) Halgren, T. A. Merck Molecular Force Field. II. MMFF94 van Der Waals and Electrostatic Parameters for Intermolecular Interactions. J. Comput. Chem. 1996, 17 (5–6), 520–552. 10.1002/(SICI)1096-987X(199604)17:5/6<520::AID-JCC2>3.0.CO;2-W.

(48) Halgren, T. A. Merck Molecular Force Field. III. Molecular Geometries and Vibrational Frequencies for MMFF94. J. Comput. Chem. 1996, 17 (5–6), 553–586. 10.1002/(SICI)1096-987X(199604)17:5/6<553::AID-JCC3>3.0.CO;2-T.

(49) Halgren, T. A.; Nachbar, R. B. Merck Molecular Force Field. IV. Conformational Energies and Geometries for MMFF94. J. Comput. Chem. 1996, 17 (5–6), 587–615. 10.1002/(SICI)1096-987X(199604)17:5/6<587::AID-JCC4>3.0.CO;2-Q.

(50) Halgren, T. A. Merck Molecular Force Field. V. Extension of MMFF94 Using Experimental Data, Additional Computational Data, and Empirical Rules. J. Comput. Chem. 1996, 17 (5–6), 616–641. 10.1002/(SICI)1096-987X(199604)17:5/6<616::AID-JCC5>3.0.CO;2-X.

(51) MacKerell, A. D.; Bashford, D.; Bellott, M.; Dunbrack, R. L.; Evanseck, J. D.; Field, M. J.; Fischer, S.; Gao, J.; Guo, H.; Ha, S.; Joseph-McCarthy, D.; Kuchnir, L.; Kuczera, K.; Lau, F. T.; Mattos, C.; Michnick, S.; Ngo, T.; Nguyen, D. T.; Prodhom, B.; Reiher, W. E.; Roux, B.; Schlenkrich, M.; Smith, J. C.; Stote, R.; Straub, J.; Watanabe, M.; Wiórkiewicz-Kuczera, J.; Yin, D.; Karplus, M. All-Atom Empirical Potential for Molecular Modeling and Dynamics Studies of Proteins. J. Phys. Chem. B 1998, 102 (18), 3586–3616. 10.1021/jp973084f.

(52) Humphrey, W.; Dalke, A.; Schulten, K. VMD: Visual Molecular Dynamics. J. Mol. Graph. 1996, 14 (1), 33–38. 10.1016/0263-7855(96)00018-5.

(53) Best, R. B.; Zhu, X.; Shim, J.; Lopes, P. E. M.; Mittal, J.; Feig, M.; MacKerell, A. D. Optimization of the Additive CHARMM All-Atom Protein Force Field Targeting Improved Sampling of the Backbone ϕ, ψ and Side-Chain χ _1_ and χ _2_ Dihedral Angles. J. Chem. Theory Comput. 2012, 8 (9), 3257–3273. 10.1021/ct300400x.

(54) Vanommeslaeghe, K.; Hatcher, E.; Acharya, C.; Kundu, S.; Zhong, S.; Shim, J.; Darian, E.; Guvench, O.; Lopes, P.; Vorobyov, I.; Mackerell, A. D. CHARMM General Force Field: A Force Field for DrugClike Molecules Compatible with the CHARMM AllCatom Additive Biological Force Fields. J. Comput. Chem. 2010, 31 (4), 671–690. 10.1002/jcc.21367.

(55) Vanommeslaeghe, K.; Raman, E. P.; MacKerell, A. D. Automation of the CHARMM General Force Field (CGenFF) II: Assignment of Bonded Parameters and Partial Atomic Charges. J. Chem. Inf. Model. 2012, 52 (12), 3155–3168. 10.1021/ci3003649.

(56) Soteras Gutiérrez, I.; Lin, F.-Y.; Vanommeslaeghe, K.; Lemkul, J. A.; Armacost, K. A.; Brooks, C. L.; MacKerell, A. D. Parametrization of Halogen Bonds in the CHARMM General Force Field: Improved Treatment of Ligand–Protein Interactions. Bioorg. Med. Chem. 2016, 24 (20), 4812–4825. 10.1016/j.bmc.2016.06.034.

(57) Kim, S.; Lee, J.; Jo, S.; Brooks, C. L.; Lee, H. S.; Im, W. CHARMM-GUI Ligand Reader and Modeler for CHARMM Force Field Generation of Small Molecules: CHARMM-GUI Ligand Reader and Modeler for CHARMM Force Field Generation of Small Molecules. J. Comput. Chem. 2017, 38 (21), 1879–1886. 10.1002/jcc.24829.

(58) Brooks, B. R.; Brooks, C. L.; Mackerell, A. D.; Nilsson, L.; Petrella, R. J.; Roux, B.; Won, Y.; Archontis, G.; Bartels, C.; Boresch, S.; Caflisch, A.; Caves, L.; Cui, Q.; Dinner, A. R.; Feig, M.; Fischer, S.; Gao, J.; Hodoscek, M.; Im, W.; Kuczera, K.; Lazaridis, T.; Ma, J.; Ovchinnikov, V.; Paci, E.; Pastor, R. W.; Post, C. B.; Pu, J. Z.; Schaefer, M.; Tidor, B.; Venable, R. M.; Woodcock, H. L.; Wu, X.; Yang, W.; York, D. M.; Karplus, M. CHARMM: The Biomolecular Simulation Program. J. Comput. Chem. 2009, 30 (10), 1545–1614. 10.1002/jcc.21287.

(59) Lee, J.; Cheng, X.; Swails, J. M.; Yeom, M. S.; Eastman, P. K.; Lemkul, J. A.; Wei, S.; Buckner, J.; Jeong, J. C.; Qi, Y.; Jo, S.; Pande, V. S.; Case, D. A.; Brooks, C. L.; MacKerell, A. D.; Klauda, J. B.; Im, W. CHARMM-GUI Input Generator for NAMD, GROMACS, AMBER, OpenMM, and CHARMM/OpenMM Simulations Using the CHARMM36 Additive Force Field. J. Chem. Theory Comput. 2016, 12 (1), 405–413. 10.1021/acs.jctc.5b00935.

(60) Phillips, J. C.; Hardy, D. J.; Maia, J. D. C.; Stone, J. E.; Ribeiro, J. V.; Bernardi, R. C.; Buch, R.; Fiorin, G.; Hénin, J.; Jiang, W.; McGreevy, R.; Melo, M. C. R.; Radak, B. K.; Skeel, R. D.; Singharoy, A.; Wang, Y.; Roux, B.; Aksimentiev, A.; Luthey-Schulten, Z.; Kalé, L. V.; Schulten, K.; Chipot, C.; Tajkhorshid, E. Scalable Molecular Dynamics on CPU and GPU Architectures with NAMD. J. Chem. Phys. 2020, 153 (4), 044130. 10.1063/5.0014475.

(61) Darden, T.; York, D.; Pedersen, L. Particle Mesh Ewald: An NClog(N) Method for Ewald Sums in Large Systems. J. Chem. Phys. 1993, 98 (12), 10089–10092. 10.1063/1.464397.

(62) Andersen, H. C. Rattle: A “Velocity” Version of the Shake Algorithm for Molecular Dynamics Calculations. J. Comput. Phys. 1983, 52 (1), 24–34. 10.1016/0021-9991(83)90014-1.

(63) Miyamoto, S.; Kollman, P. A. Settle: An Analytical Version of the SHAKE and RATTLE Algorithm for Rigid Water Models. J. Comput. Chem. 1992, 13 (8), 952–962. 10.1002/jcc.540130805.

(64) Gowers, R.; Linke, M.; Barnoud, J.; Reddy, T.; Melo, M.; Seyler, S.; Domański, J.; Dotson, D.; Buchoux, S.; Kenney, I.; Beckstein, O. MDAnalysis: A Python Package for the Rapid Analysis of Molecular Dynamics Simulations; Austin, Texas, 2016; pp 98–105. 10.25080/Majora-629e541a-00e.

(65) MichaudCAgrawal, N.; Denning, E. J.; Woolf, T. B.; Beckstein, O. MDAnalysis: A Toolkit for the Analysis of Molecular Dynamics Simulations. J. Comput. Chem. 2011, 32 (10), 2319–2327. 10.1002/jcc.21787.

(66) Goddard, T. D.; Huang, C. C.; Meng, E. C.; Pettersen, E. F.; Couch, G. S.; Morris, J. H.; Ferrin, T. E. UCSF ChimeraX: Meeting Modern Challenges in Visualization and Analysis. Protein Sci. 2018, 27 (1), 14–25. 10.1002/pro.3235.

(67) Meng, E. C.; Goddard, T. D.; Pettersen, E. F.; Couch, G. S.; Pearson, Z. J.; Morris, J. H.; Ferrin, T. E. UCSF CHIMERAXC: Tools for Structure Building and Analysis. Protein Sci. 2023, 32 (11), e4792. 10.1002/pro.4792.

(68) Pettersen, E. F.; Goddard, T. D.; Huang, C. C.; Meng, E. C.; Couch, G. S.; Croll, T. I.; Morris, J. H.; Ferrin, T. E. UCSF CHIMERAXC: Structure Visualization for Researchers, Educators, and Developers. Protein Sci. 2021, 30 (1), 70–82. 10.1002/pro.3943.

(69) Wallace, A. C.; Laskowski, R. A.; Thornton, J. M. LIGPLOT: A Program to Generate Schematic Diagrams of Protein-Ligand Interactions. Protein Eng. 1995, 8 (2), 127–134. 10.1093/protein/8.2.127.

(70) Laskowski, R. A.; Swindells, M. B. LigPlot+: Multiple Ligand-Protein Interaction Diagrams for Drug Discovery. J. Chem. Inf. Model. 2011, 51 (10), 2778–2786. 10.1021/ci200227u.

(71) Gao, X.; Shi, Z.; Zhou, R. FEbuilder: A Comprehensive Webserver to Streamline FEP Simulation Setup in Drug Discovery, 2024. 10.5281/zenodo.14059079.

(72) Kwiatkowski, N.; Zhang, T.; Rahl, P. B.; Abraham, B. J.; Reddy, J.; Ficarro, S. B.; Dastur, A.; Amzallag, A.; Ramaswamy, S.; Tesar, B.; Jenkins, C. E.; Hannett, N. M.; McMillin, D.; Sanda, T.; Sim, T.; Kim, N. D.; Look, T.; Mitsiades, C. S.; Weng, A. P.; Brown, J. R.; Benes, C. H.; Marto, J. A.; Young, R. A.; Gray, N. S. Targeting Transcription Regulation in Cancer with a Covalent CDK7 Inhibitor. Nature 2014, 511 (7511), 616–620. 10.1038/nature13393.

(73) Krenske, E. H.; Petter, R. C.; Houk, K. N. Kinetics and Thermodynamics of Reversible Thiol Additions to Mono- and Diactivated Michael Acceptors: Implications for the Design of Drugs That Bind Covalently to Cysteines. J. Org. Chem. 2016, 81 (23), 11726–11733. 10.1021/acs.joc.6b02188.

