## Supporting Information for "A Robust Computational Framework for the Optimization of CDK7 Inhibitors as Promising Cancer Therapy"

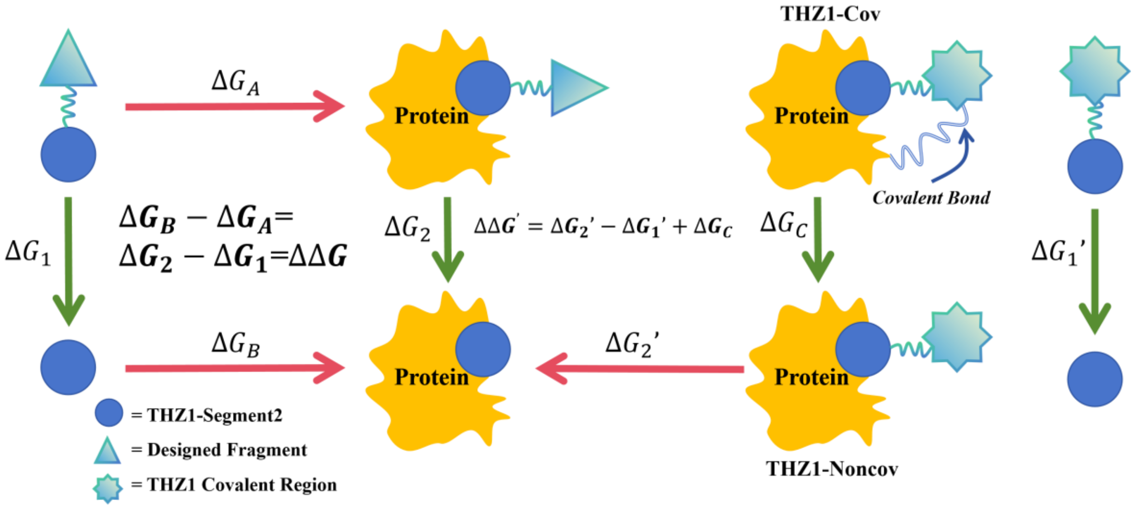


**Figure S1.** Free Energy Perturbation (FEP) workflow. The left panel illustrates the classical FEP thermodynamic cycle, where all newly designed molecules are transformed to Segment2. The right panel depicts calculating the relative binding free energy (ΔΔG) between THZ1-Cov and Segment2. The free energy change of the reversed Michael addition reaction ($\Delta\Delta G_C$) is obtained from DFT calculations1. Subsequently, THZ1-Noncov is transformed to Segment2 via standard FEP ($\Delta G_2' - \Delta G_1'$). We can compare $\Delta\Delta G = \Delta G_B - \Delta G_A = \Delta G_2 - \Delta G_1$ and $\Delta\Delta G' = \Delta G_2' - \Delta G_1' + \Delta G_C$ to estimate whether the newly designed molecules are more favorable than covalent THZ1.

**Table S1. Structures and SMILES notation of DiffLinker-generated candidate CDK7 inhibitors (CDIs) for molecular dynamics (MD) simulation.**

| Candidates | Structure | SMILES |
|---|---|---|
| CDI1 | 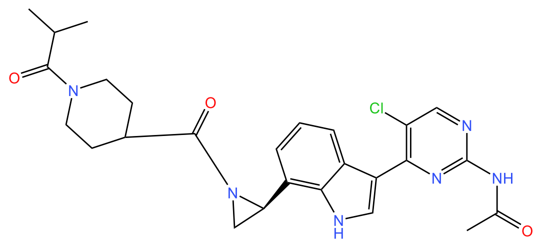 | Clc1cnc(NC(=O)C)nc1c1c[nH]c2c(cccc12)[C@@H]1N(C(=O)C2CCN(CC2)C(=O)C(C)C)C1 |
| CDI2 | 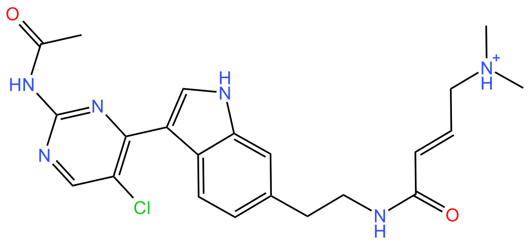 | Clc1cnc(NC(=O)C)nc1c1c[nH]c2cc(ccc12)CCNC(=O)/C=C/C[NH+](C)C |
| CDI3 | 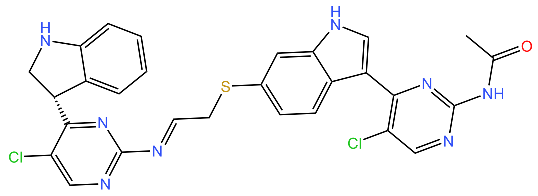 | Clc1cnc(NC(=O)C)nc1c1c[nH]c2cc(ccc12)SC/C=N/c1ncc(c(n1)[C@@H]1CNc2ccccc12)Cl |
| CDI4 | 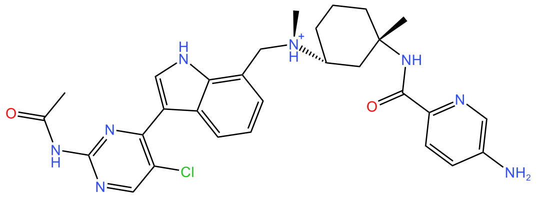 | Clc1cnc(NC(=O)C)nc1c1c[nH]c2c(cccc12)C[N@H+]([C@@H]1CCC[C@@](C1)(NC(=O)c1ncc(cc1)N)C)C |

CDI5

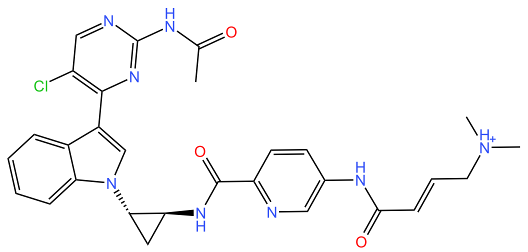

Clc1cnc(NC(=O)C)
nc1c1cn(c2ccccc12)
[C@H]1C[C@@H]
1NC(=O)c1ncc(cc1)
NC(=O)/C=C/C[NH
+](C)C

CDI6

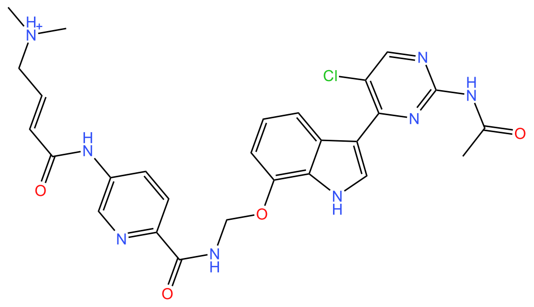

Clc1cnc(NC(=O)C)
nc1c1c[nH]c2c(cccc
12)OCNC(=O)c1ncc
(cc1)NC(=O)/C=C/
C[NH+](C)C

CDI7

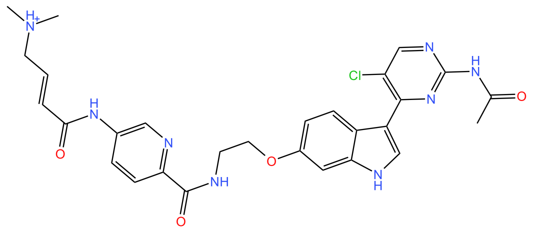

Clc1cnc(NC(=O)C)
nc1c1c[nH]c2cc(ccc
12)OCCNC(=O)c1n
cc(cc1)NC(=O)/C=
C/C[NH+](C)C

CDI8

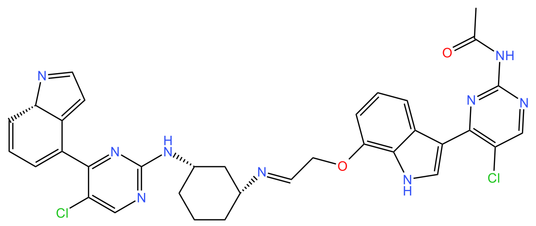

Clc1cnc(NC(=O)C)
nc1c1c[nH]c2c(cccc
12)OC/C=N/[C@@
H]1CCC[C@@H](
C1)Nc1ncc(c(n1)C1
=CC=C[C@H]2C1=
CC=N2)Cl

CDI9

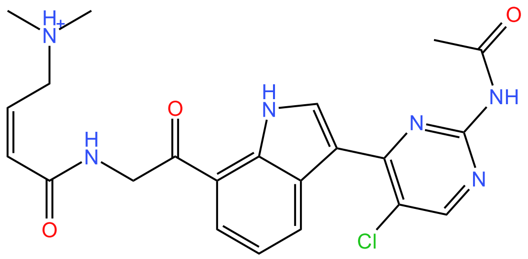

Clc1cnc(NC(=O)C)
nc1c1c[nH]c2c(cccc
12)C(=O)CNC(=O)/
C=C\C[NH+](C)C

CDI10

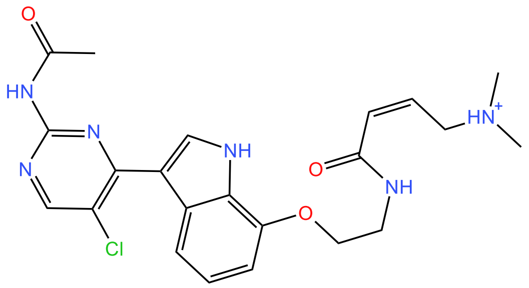

Clc1cnc(NC(=O)C)
nc1c1c[nH]c2c(cccc
12)OCCNC(=O)/C=
C\C[NH+](C)C

CDI11

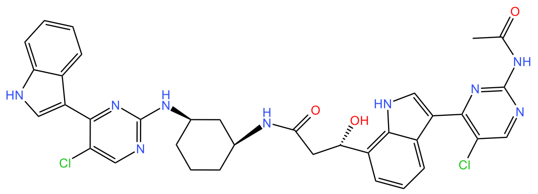

Clc1cnc(NC(=O)C)
nc1c1c[nH]c2c(cccc
12)[C@H](CC(=O)
N[C@H]1CCC[C@
H](C1)Nc1ncc(c(n1)
c1c[nH]c2ccccc12)
Cl)O

CDI12

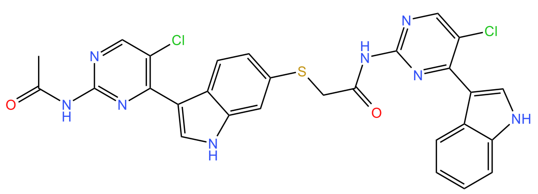

Clc1cnc(NC(=O)C)
nc1c1c[nH]c2cc(ccc
12)SCC(=O)Nc1ncc
(c(n1)c1c[nH]c2ccc
cc12)Cl

CDI13

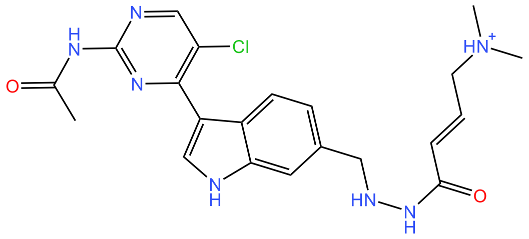

Clc1cnc(NC(=O)C)
nc1c1c[nH]c2cc(ccc
12)CNNC(=O)/C=C
/C[NH+](C)C

CDI14

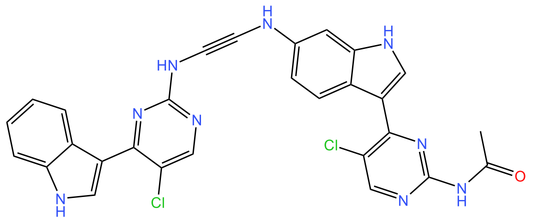

Clc1cnc(NC(=O)C)
nc1c1c[nH]c2cc(ccc
12)NC#CNc1ncc(c(
n1)c1c[nH]c2ccccc1
2)Cl

| | | |
|---|---|---|
| CDI15 | 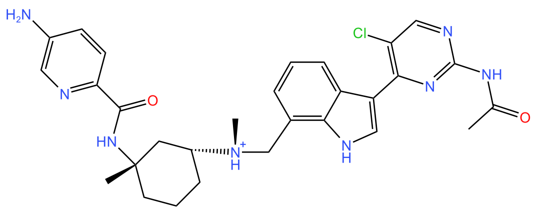 | Clc1cnc(NC(=O)C)nc1c1c[nH]c2c(cccc12)C[N@H+]([C@@H]1CCC[C@@](C1)(NC(=O)c1ncc(cc1)N)C)C |
| CDI16 | 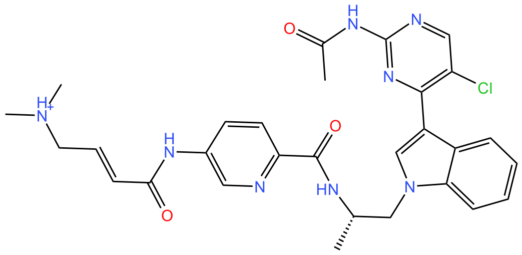 | Clc1cnc(NC(=O)C)nc1c1cn(c2ccccc12)C[C@@H](NC(=O)c1ncc(cc1)NC(=O)/C=C/C[NH+](C)C)C |
| CDI18 | 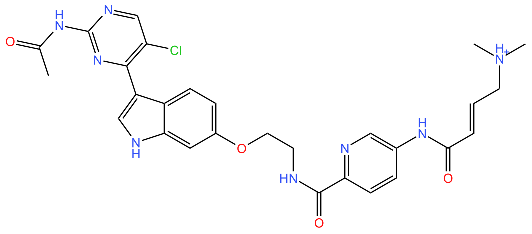 | Clc1cnc(NC(=O)C)nc1c1c[nH]c2cc(ccc12)OCCNC(=O)c1ncc(cc1)NC(=O)/C=C/C[NH+](C)C |
| CDI19 | 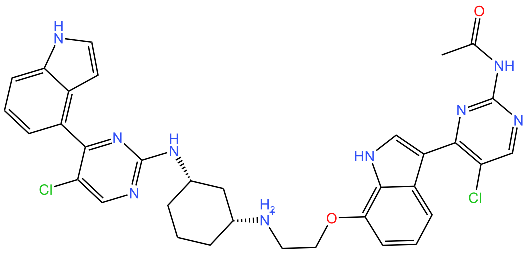 | Clc1cnc(NC(=O)C)nc1c1c[nH]c2c(cccc12)OCC[NH2+][C@@H]1CCC[C@@H](C1)Nc1ncc(c(n1)c1c2cc[nH]c2ccc1)Cl |

CDI20

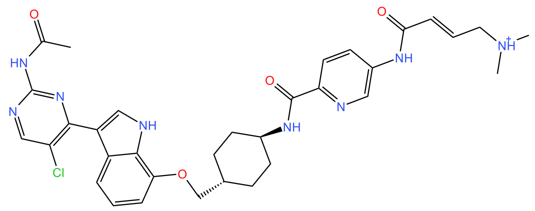

Clc1cnc(NC(=O)C)nc1c1c[nH]c2c(cccc12)OC[C@@H]1CC[C@H](CC1)NC(=O)c1ncc(cc1)NC(=O)/C=C/C[NH+](C)C

CDI21

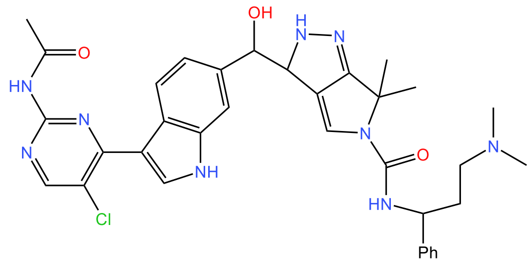

Clc1cnc(NC(=O)C)nc1c1c[nH]c2cc(ccc12)[C@@H]([C@H]1C2=CN(C(C)(C2=NN1)C)C(=O)N[C@H](c1ccccc1)C[NH+](C)C)O

CDI22

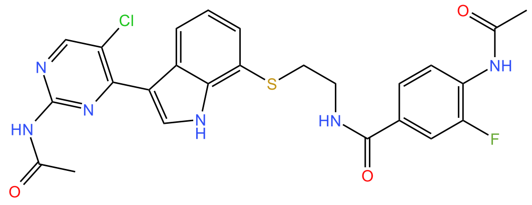

Clc1cnc(NC(=O)C)nc1c1c[nH]c2c(cccc12)SCCNC(=O)c1cc(c(cc1)NC(=O)C)F

CDI23

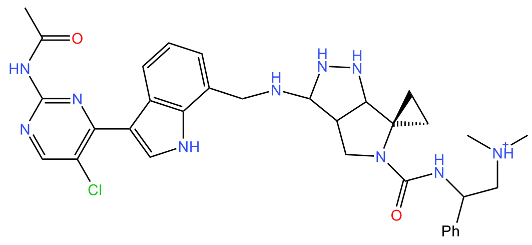

Clc1cnc(NC(=O)C)nc1c1c[nH]c2c(cccc12)CNC1=[NH]NC2=C1CN(C(=O)N[C

| | | |
|---|---|---|
| | | @H](c1ccccc1)C[NH+](C)C)C12CC1 |
| CDI24 | 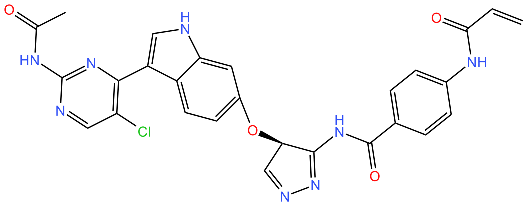 | Clc1cnc(NC(=O)C)nc1c1c[nH]c2cc(ccc12)O[C@@H]1C=NN=C1NC(=O)c1ccc(cc1)NC(=O)C=C |
| CDI25 | 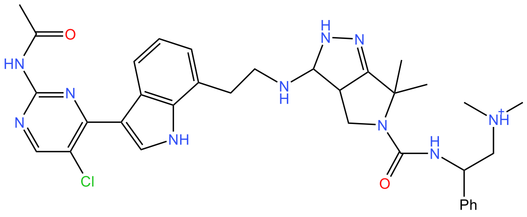 | Clc1cnc(NC(=O)C)nc1c1c[nH]c2c(cccc12)CCNC1=[NH]N=C2[C@H]1CN(C2(C)C)C(=O)N[C@@H](c1ccccc1)C[NH+](C)C |
| CDI26 | 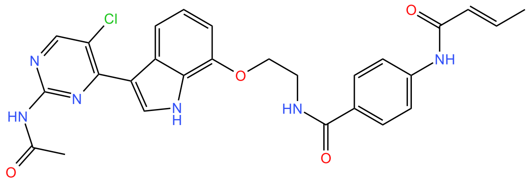 | Clc1cnc(NC(=O)C)nc1c1c[nH]c2c(cccc12)OCCNC(=O)c1ccc(cc1)NC(=O)/C=C/C |
| CDI27 | 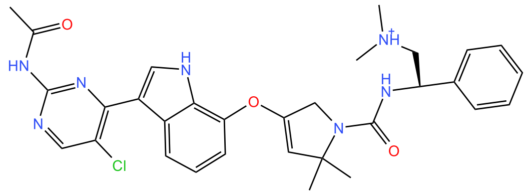 | Clc1cnc(NC(=O)C)nc1c1c[nH]c2c(cccc12)OC1=CC(N(C1)C(=O)N[C@@H](c1ccccc1)C[NH+](C)C)(C)C |

CDI28

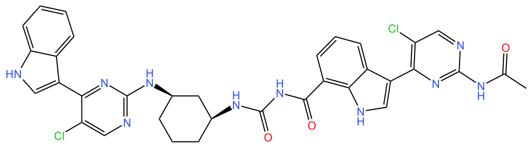


Clc1cnc(NC(=O)C)nc1c1c[nH]c2c(cccc12)C(=O)NC(=O)N[C@H]1CCC[C@H](C1)Nc1ncc(c(n1)c1c[nH]c2ccccc12)Cl

CDI29

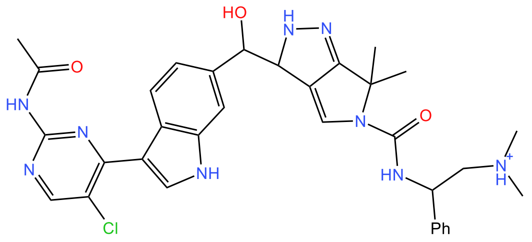


Clc1cnc(NC(=O)C)nc1c1c[nH]c2cc(ccc12)C(=O)[C@H]1C2=CN(C(C)(C2=NN1)C)C(=O)N[C@@H](c1ccccc1)C[NH+](C)C

CDI30

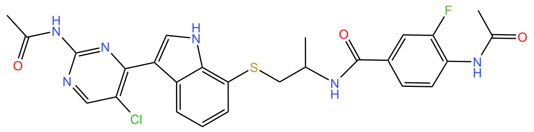


N(C(=O)c1cc(c(cc1)NC(=O)C)F)C(CSc1c2[nH]cc(c3c(Cl)cnc(NC(=O)C)n3)c2ccc1)C

CDI31

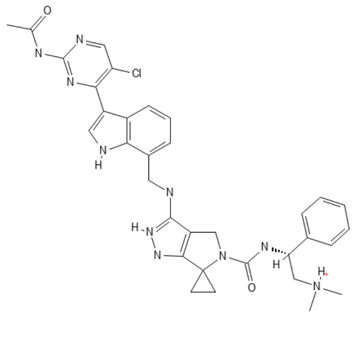


Clc1cnc(NC(=O)C)nc1c1c[nH]c2c(cccc12)CNC1=[NH]NC2=C1CN(C(=O)N[C@H](c1ccccc1)C[NH+](C)C)C12CC1

| | | |
|---|---|---|
| CDI32 | 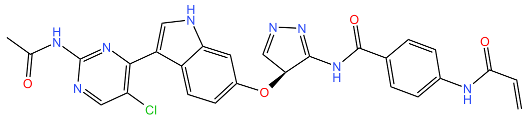 | Clc1cnc(NC(=O)C)nc1c1c[nH]c2cc(ccc12)O[C@H]1C=NN=C1NC(=O)c1ccc(cc1)NC(=O)C=C |
| CDI33 | 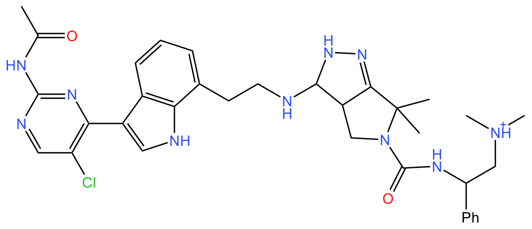 | Clc1cnc(NC(=O)C)nc1c1c[nH]c2c(cccc12)CCNC1=[NH]N=C2[C@H]1CN(C2(C)C)C(=O)N[C@H](c1ccccc1)C[NH+](C)C |
| CDI34 | 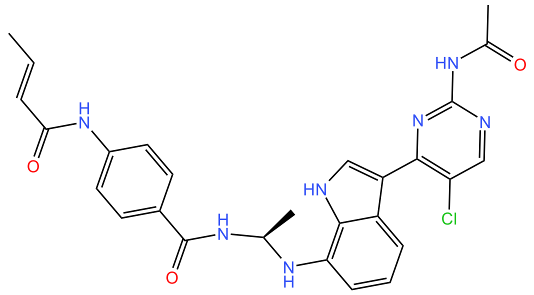 | Clc1cnc(NC(=O)C)nc1c1c[nH]c2c(cccc12)N[C@@H](NC(=O)c1ccc(cc1)NC(=O)/C=C/C)C |
| CDI35 | 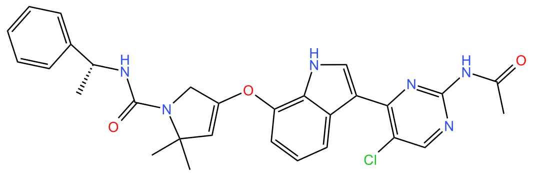 | Clc1cnc(NC(=O)C)nc1c1c[nH]c2c(cccc12)OC1=CC(N(C1)C(=O)N[C@@H](c1ccccc1)C)(C)C |

CDI36

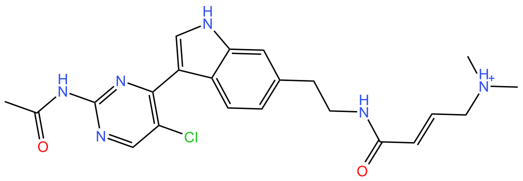

Clc1cnc(NC(=O)C)nc1c1c[nH]c2cc(ccc12)CCNC(=O)/C=C/C[NH+](C)C

CDI37

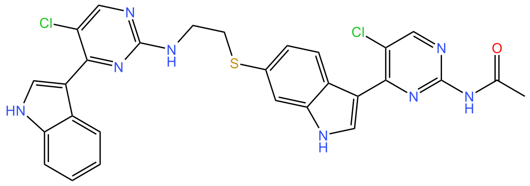

Clc1cnc(NC(=O)C)nc1c1c[nH]c2cc(ccc12)SCCNc1ncc(c(n1)c1c[nH]c2ccccc12)Cl

CDI38

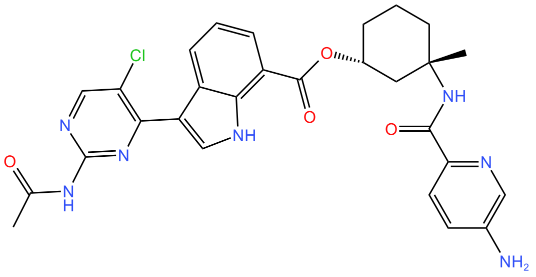

Clc1cnc(NC(=O)C)nc1c1c[nH]c2c(cccc12)C(=O)O[C@@H]1CCC[C@@](C1)(NC(=O)c1ncc(cc1)N)C

CDI39

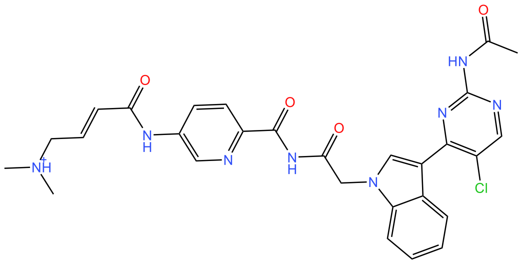

Clc1cnc(NC(=O)C)nc1c1cn(c2ccccc12)CC(=O)NC(=O)c1ncc(cc1)NC(=O)/C=C/C[NH+](C)C

CDI41

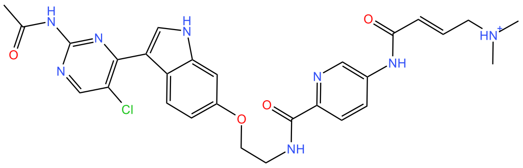

Clc1cnc(NC(=O)C)nc1c1c[nH]c2cc(ccc12)OCCNC(=O)c1ncc(cc1)NC(=O)/C=C/C[NH+](C)C

| | | |
|---|---|---|
| CDI42 | 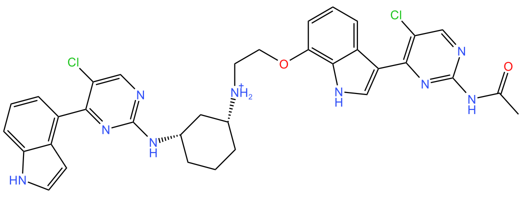 | Clc1cnc(NC(=O)C)nc1c1c[nH]c2c(cccc12)OCC[NH2+][C@@H]1CCC[C@@H](C1)Nc1ncc(c(n1)c1c2cc[nH]c2ccc1)Cl |
| CDI43 | 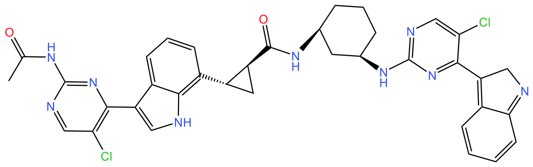 | Clc1cnc(NC(=O)C)nc1c1c[nH]c2c(cccc12)[C@@H]1[C@@H](C(=O)N[C@H]2CCC[C@H](C2)Nc2ncc(c(n2)C2=c3c(=NC2)cccc3)Cl)C1 |
| CDI45 | 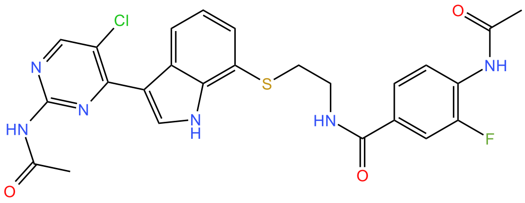 | Clc1cnc(NC(=O)C)nc1c1c[nH]c2c(cccc12)SCCNC(=O)c1cc(c(cc1)NC(=O)C)F |
| CDI46 | 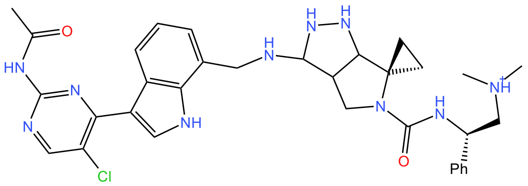 | Clc1cnc(NC(=O)C)nc1c1c[nH]c2c(cccc12)CNC1=[NH]NC2=C1CN(C(=O)N[C@H](c1ccccc1)C[NH+](C)C)C12CC1 |

| Compound | Structure | SMILES |
|---|---|---|
| CDI47 | 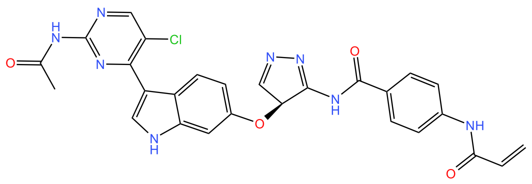 | Clc1cnc(NC(=O)C)nc1c1c[nH]c2cc(ccc12)O[C@H]1C=NN=C1NC(=O)c1ccc(cc1)NC(=O)C=C |
| CDI48 | 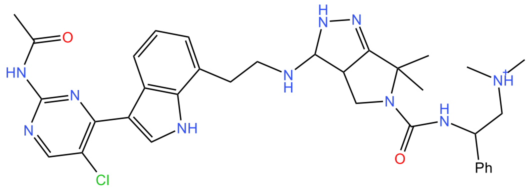 | Clc1cnc(NC(=O)C)nc1c1c[nH]c2c(cccc12)CCNC1=[NH]N=C2[C@H]1CN(C2(C)C)C(=O)N[C@@H](c1ccccc1)C[NH+](C)C |
| CDI49 | 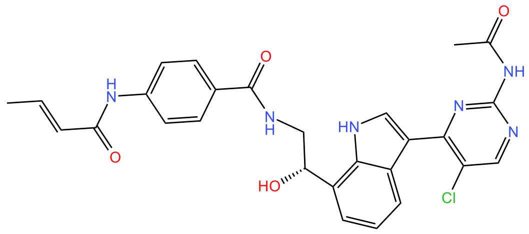 | Clc1cnc(NC(=O)C)nc1c1c[nH]c2c(cccc12)[C@@H](CNC(=O)c1ccc(cc1)NC(=O)/C=C/C)O |
| CDI50 | 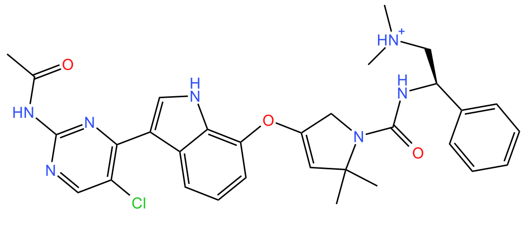 | Clc1cnc(NC(=O)C)nc1c1c[nH]c2c(cccc12)OC1=CC(N(C1)C(=O)N[C@H](c1ccccc1)C[NH+](C)C)(C)C |

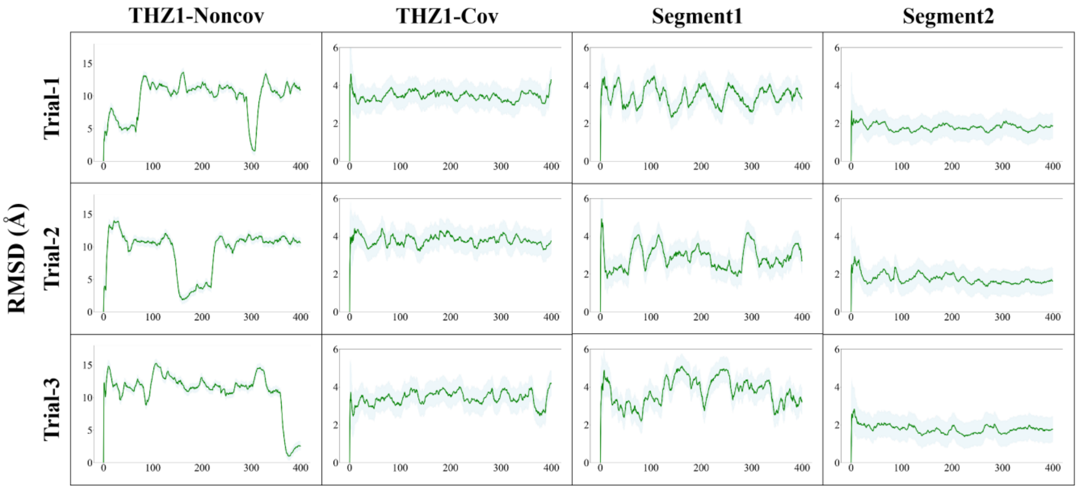


**Figure S2.** Root mean squared deviation (RMSD) of THZ1-Noncov, THZ1-Cov, Segment1, and Segment2 during MD simulation. Each molecule was subjected to three parallel MD trials. The covalent bond helps THZ1-Cov remain stable within the pocket. Conversely, without a covalent bond, THZ1-Noncov becomes highly unstable. Both Segment1 and Segment2 are stable.

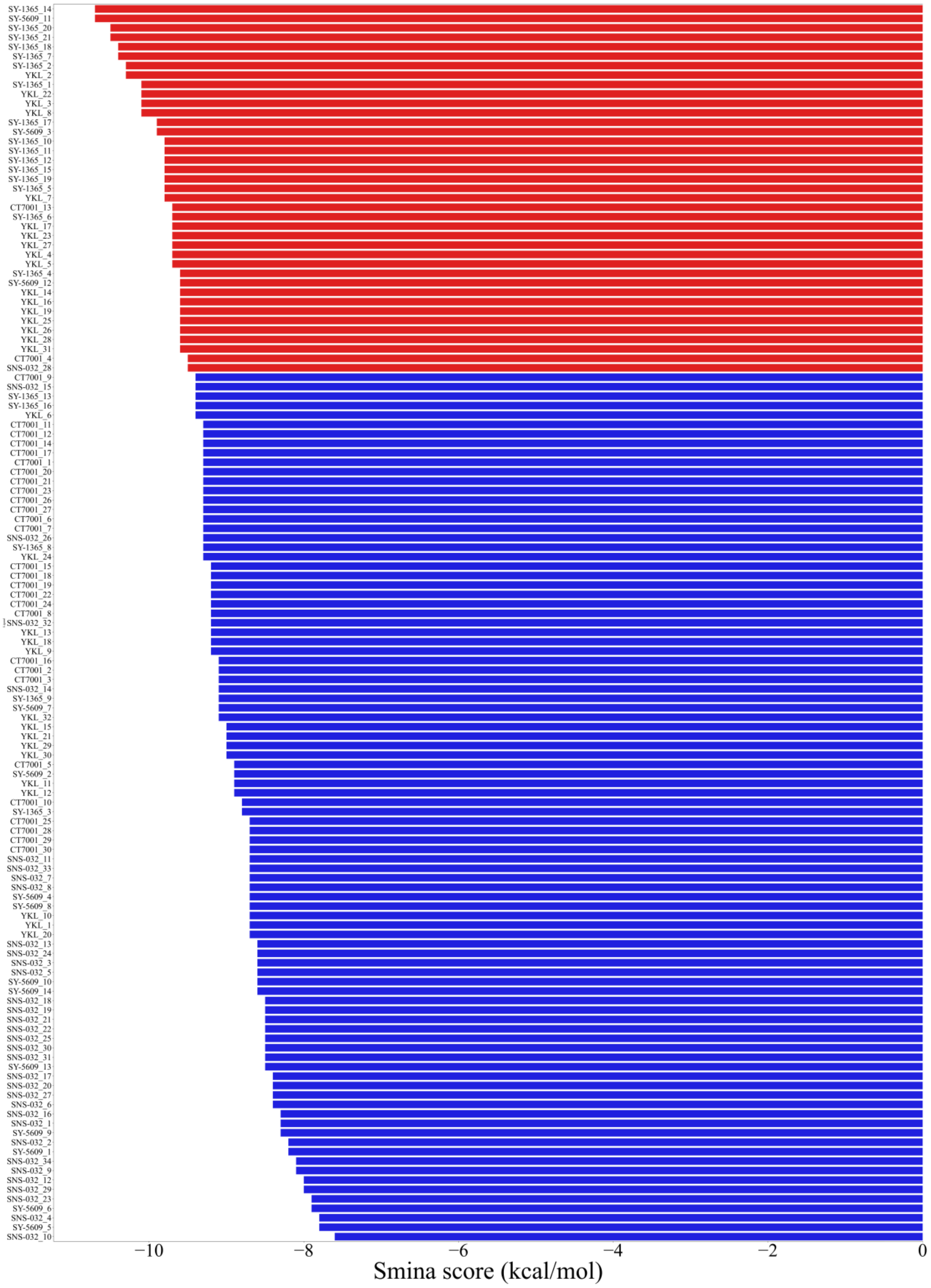


**Figure S3.** Sorted smina scores of eight known CDK7 inhibitors and their derivatives in the first round of docking. Lower smina scores (kcal/mol) indicate stronger binding affinity. The red bars indicate small molecules with a docking score less than 1.2 kcal/mol higher than the best ligand, indicating favorable binding affinity. These molecules were retained as candidate molecules for further screening. The blue bars represent small molecules with less favorable scores, which were discarded after this round.

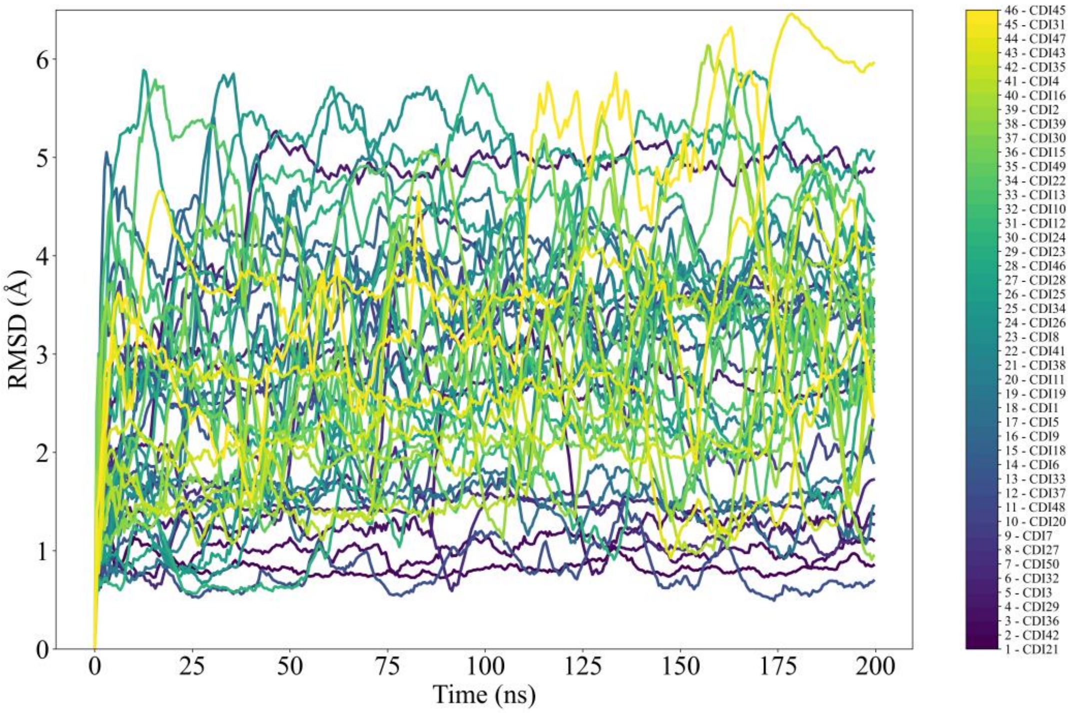


**Figure S4.** Stability ranking of dual-site inhibitors over 200 ns MD simulations. The RMSD values for each ligand were monitored to evaluate conformational stability. Molecules are ranked by the standard deviation of RMSD calculated over the last 50 ns of simulation, with lower deviations (or darker color) indicating higher stability.

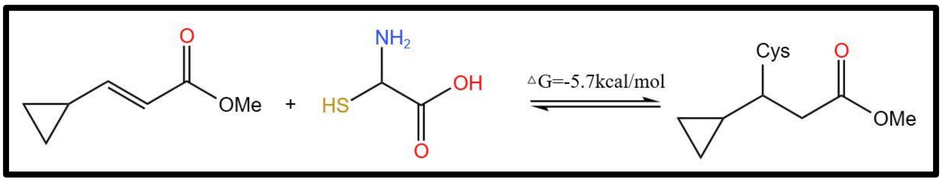


**Figure S5.** Free energy change for cysteine binding via Michael addition. The free energy change (ΔG) for the covalent bond formation between cysteine and 'molecule 18' (from Krenske et al. in ref 1) was calculated as -5.7 kcal/mol[1], as determined using density functional theory (DFT) with the M06-2X functional, a 6-311+G(d,p) basis set, and the CPCM solvent model to simulate an aqueous environment.

### Reference

(1) Krenske, E. H.; Petter, R. C.; Houk, K. N. Kinetics and Thermodynamics of Reversible Thiol Additions to Mono- and Diactivated Michael Acceptors: Implications for the Design of Drugs That Bind Covalently to Cysteines. *J. Org. Chem.* **2016**, *81* (23), 11726–11733. https://doi.org/10.1021/acs.joc.6b02188.
