## Supplementary material for "A Robust Computational Framework for the Optimization of CDK7 Inhibitors as Promising Cancer Therapy": Cover Letter

ZHEJIANG UNIVERSITY

Qiushi Chair Professor and Director
Institute of Quantitative Biology
Zhejiang University, China


November 12th, 2024

Dear *BioRxiv* editors,

We are pleased to submit the manuscript entitled "***A Robust Computational Framework for the Optimization of CDK7 Inhibitors as Promising Cancer Therapy***" for your kind consideration of publication in *BioRxiv*. The following is a brief description of our work and its significance in the future research and development of CDK7 inhibitors.

We present a streamlined computational approach to design non-covalent CDK7 inhibitors with enhanced selectivity and reduced off-target effects. CDK7, a critical component of the CDK-activating kinase (CAK) complex, plays key roles in gene transcription and cell cycle regulation. Its overexpression in tumors, such as triple-negative breast cancer (TNBC) and lung squamous cell carcinoma (LSCC), supports their malignant phenotypes. Inhibiting CDK7 can induce apoptosis in TNBC cells and control LSCC cell proliferation and survival by reducing SOX2 expression, highlighting the therapeutic potential of CDK7 inhibitors.

Our study focuses on the ATP-binding site and active site of CDK7, utilizing molecular dynamics (MD) simulations, molecular docking, and free energy perturbation (FEP) to identify promising small-molecule inhibitors. Initial MD simulations of the THZ1-Cyclin H-CDK7 (CAK) complex revealed that THZ1-Cov (covalent form) maintains stable interactions with key residues, while THZ1-Noncov (non-covalent form) exhibits conformational fluctuations but retains stable interactions within the core pocket, particularly with residues Leu18, Val26, Phe93, Leu144, and Asp155 **(Figure R1. A-B)**. To further investigate the role of core pocket, we conducted MD simulations of two simplified THZ1 fragments—THZ1-Segment1 and THZ1-Segment2. Both fragments showed stable binding with minimal conformational changes, closely resembling the binding patterns of THZ1-Cov and THZ1-Noncov. These results indicate the importance of core pocket and suggest THZ1-Segment2 as a promising scaffold to design non-covalent inhibitors.

We identified 147 molecules from a similarity search of eight known CDK7 inhibitors in the PubChem database. These molecules were docked into a box encompassing the core pocket and adjacent pockets. The top candidates (smina scores below -9.5 kcal/mol) were selected for further analysis. From this subset, we extracted molecular fragments binding to adjacent pockets and used the deep learning program DiffLinker to generate linkers between these fragments and THZ1-Segment2, considering pocket geometry. After manual inspection, 46 candidates were identified as potential CDK7 inhibitors. We assessed the structural stability of these candidates using 200 ns MD simulations. The top 17 ligands, exhibited minimal conformational changes and were advanced to FEP calculations. We adopted -5.7 kcal/mol from previous density functional theory (DFT) calculations as the free energy change of Cys312 added to THZ1. The baseline is ΔΔG to transform THZ1-Cov to THZ1-Segment2.

FEP calculations revealed that most designed fragments have enhanced binding affinities relative to THZ1-Noncov. Notably, CDI3, CDI18, and CDI37 exhibited significantly higher ΔΔG values **(Figure R1 C)**. In the core pocket, THZ1-Segment2 forms a stable hydrogen bond with Met94 and additional interactions with Leu18, Val26, Phe93, and Leu144, as well as polar contacts with Asp92 and Asp155. These interactions resemble those in THZ1-Cov and THZ1-Noncov.

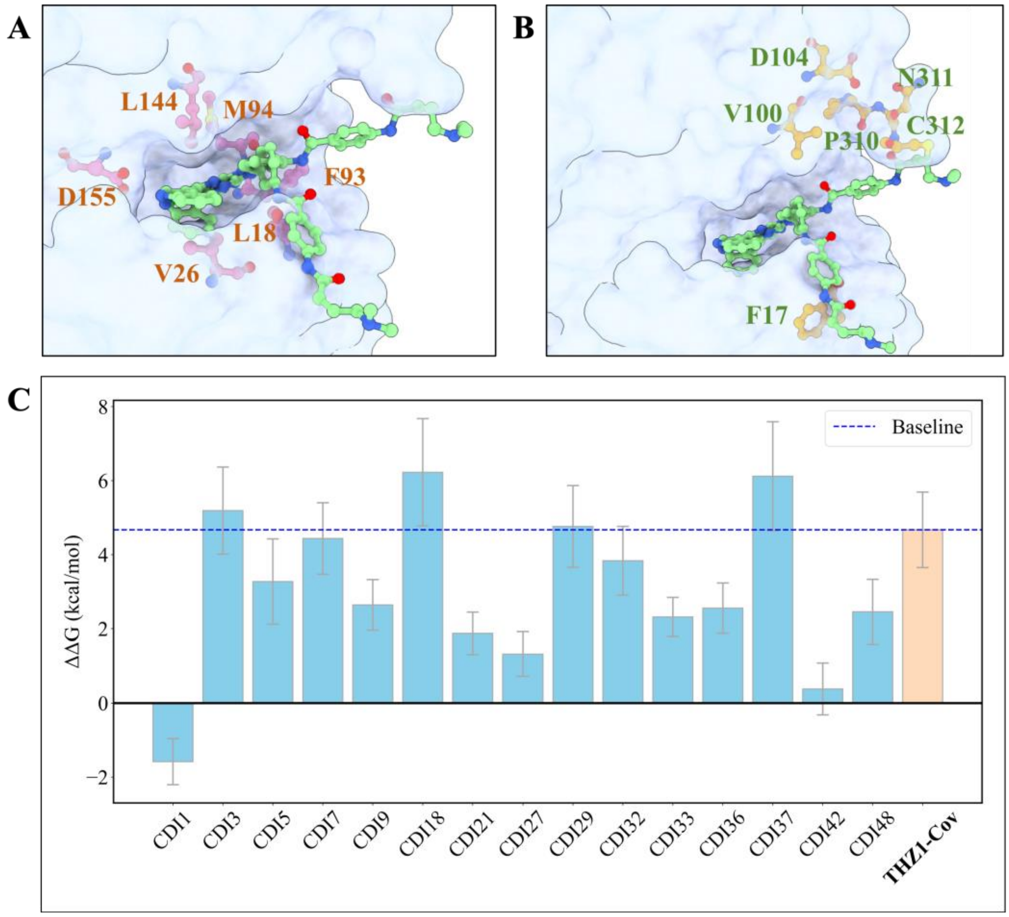


**Figure R1. Relative binding free energy and of designed CDK7 inhibitors.** (A-B) Detailed views of THZ1-Cov and THZ1-Noncov in the CDK7 pocket, demonstrating distinct binding conformations. (C) Bar plot of ΔΔG for various CDIs (blue) transforming into THZ1-Segment2. Error bars indicate standard deviations. THZ1-Cov (orange) is the baseline, accounting for covalent bond formation energy plus ΔΔG transforming THZ1-Noncov to THZ1-Segment2. Positive ΔΔG values indicate higher binding affinity, with CDI3, CDI18, and CDI37 exhibiting the strongest binding affinities.

CDI3 and CDI37 show distinct interactions in pocket 2, with CDI3 forming key contacts with residues Glu99, Lys103, and Asn141, and CDI37 engaging with a broader network including Gln22, Phe93, and Asn142. The purine group in CDI37 forms a particularly stable hydrogen bond with Asp137. CDI18, structurally similar to THZ1 with a shorter linker, primarily inserts into pocket 1, forming hydrogen bonds with Asp137 and interactions with Lys139 and Leu158, as well as electrostatic with Gln172, Val174, and π-cation interactions with Phe23.

In summary, our study provides a robust computational workflow to design non-covalent CDK7 inhibitors, offering safer alternatives and valuable insights for future kinase-targeted therapies. We are confident that our methods and findings will contribute to the design and optimization of next-generation CDK7 inhibitors, ultimately contributing to more effective and safer cancer treatments.

Thank you for considering our submission. We look forward to your feedback and are happy to provide any additional information or support that may be required.

Thanks.

Sincerely,

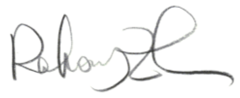

Ruhong Zhou, PhD
Zhejiang University and Columbia University
